# A genome-wide scan for correlated mutations detects macromolecular and chromatin interactions in *Arabidopsis thaliana*

**DOI:** 10.1101/279489

**Authors:** Laura Perlaza-Jiménez, Dirk Walther

## Abstract

The concept of exploiting correlated mutations has been introduced and applied successfully to identify interactions within and between biological macromolecules. Its rationale lies in the preservation of physical interactions via compensatory mutations. With the massive increase of available sequence information, approaches based on correlated mutations have regained considerable attention.

We analyzed a set of 10,707,430 single nucleotide polymorphisms detected in 1,135 accessions of the plant *Arabidopsis thaliana*. To measure their covariance and to reveal the global genome-wide sequence correlation structure of the Arabidopsis genome, the adjusted mutual information has been estimated for each possible pair of polymorphic sites. We developed a series of filtering steps to account for genetic linkage and lineage relations between Arabidopsis accessions, as well as transitive covariance as possible confounding factors. We show that upon appropriate filtering, correlated mutations prove indeed informative with regard to molecular interactions, and furthermore, appear to reflect on chromosomal interactions.

Our study demonstrates that the concept of correlated mutations can also be applied successfully to within-species sequence variation and establishes a promising approach to help unravel the complex molecular interactions in *A. thaliana* and other species with broad sequence information.

## Introduction

At the molecular level, biological functions result from complex and dynamic interactions between individual molecules (e.g. protein-protein, miRNA-mRNAs, DNA-DNA, DNA-RNA). With the great variety of possible molecular interactions, the methods to experimentally detect and computationally predict them are also very diverse (e.g.(1–3)). Despite tremendous progress, obtaining a comprehensive view of interactions of all genome-encoded molecules and genomic elements remains highly challenging. However, independent of the type of interaction, there is one assumption in common. Molecules “communicate” between each other through structure-mediated physical interactions. Thus, the information that defines the structure of molecules, i.e. their sequences or molecular makeup, should contain information about their molecular interactions. Given the challenges associated with predicting structures and their interactions based on favorable – in the sense of low free energy - physical interactions, statistical approaches have been developed that harness the wealth of genome sequence information to inform on molecular interactions. Such statistical approaches exploit covariation at different sites detected in alignments of genomic/protein/RNA sequences (4–10). Their rationale relies on the concept of compensatory mutations. As one interaction partner is mutated, leading to a change of structure or interaction surface potential, natural selection will favor mutations in the cognate molecule that compensate for this change in order to preserve the interaction. Thus, correlated mutations can be taken as indicators of possible molecular interactions and can be detected by performing covariation analyses of sets of aligned sequences.

In the plant *Arabidopsis thaliana*, the application of the concept of covariance to deduce structural and functional interactions has been demonstrated successfully for the prediction of structure-function relationships of the pentatricopeptide repeat proteins (PPR). *In silico* and experimental evidence has demonstrated that covariance in genomic sequences of orthologous protein-RNA pairs allow detecting protein-RNA recognition events (5). This covariance was identified by finding correlated mutations in the alignments between PPR binding factors and their RNA targets from different species (5–7, 11, 12). Correlated mutations have also been used successfully to identify DNA-binding domains and sites of transcription factors, such as MerR-family proteins (13). Extended to whole genomes, this concept was used in the Drosophila genus (14). In this study, investigators searched for long-range covariation clusters on a genomic scale based on genome-triple “fingerprints” and reported that compensatory mutations suggest long range interactions between exons of mRNAs and also between noncoding RNAs.

Although the rationale of using correlated mutations to inform on molecular interactions seems very plausible and straightforward, it has also been shown that covarying sites do not necessarily correspond to sites that are spatially interacting (4–10, 15). Phylogenetic bias, transitivity effects, passenger mutations are some of the reasons why the detected covariance can be misleading (4). However, several approaches have been devised that circumvent these methodological problems. Maximum-Entropy probability models (MEPM) have been developed to predict the 3D-structure of proteins and RNA-molecules as well as protein-protein and protein-RNA interactions by using global statistical models of covariation over deep alignments of gene families (8–10). MEPMs were adapted for each case of interaction or genomic element, which allowed avoiding the transitivity effect and phylogenetic sampling bias depending on the genomic element of study (10). All these approaches have been implemented using alignments of sequences from diverse species, avoiding, or at least reducing, any phylogenetic bias. By contrast, addressing compensatory mutations to detect functional associations in closely related genomes remains a challenge.

In this study, we set out to exploit the recently released genome sequence information of 1,135 genomes of *Arabidopsis thaliana* accessions (16). Thus, we focused on intra-species variation with mutations represented as single nucleotide polymorphisms (SNPs). This dataset contains around 10 million polymorphic sites, and all possible pairs were tested exhaustively for correlated mutations as a means to detect functional associations. To determine the molecules involved in these functional associations, the availability of detailed genome annotations is critical. In Arabidopsis, about 77% of the genes in the genome have been annotated (17), and other genomic elements such as pseudogenes, and mobile elements have also been identified. To detect covariance, we used as a metric the Adjusted Mutual Information (AMI) between pairs of polymorphic sites (18). When working with genomes from the same species, tight covariation can also emerge from factors other than compensatory selection such as linkage, lineage (population structure), and transitivity. We devised strategies to control for these confounding factors and to enrich for correlated mutations pairs with functional relevance. We provide a genome-wide covariation statistic and report higher than expected events of correlated mutations in molecules that have been reported to interact (protein-protein, miRNA-mRNA) as well as present evidence for the validity of interactions predicted based on detected covariance. Most strikingly, we found that genomic sites reported to physically interact to form chromatin loops and long-range chromosomal contacts were enriched for correlated mutations suggesting a link between 3D-genome organization and the sequence correlation structure.

## Methods

### Polymorphic site information

We used sequencing data and variant calls as available as part of the Arabidopsis 1001 genome project including genomic sequences of 1,135 *A. thaliana* accessions (16). We considered the nuclear genome only (chromosomes 1–5) and required polymorphic sites to be biallelic and excluding any insertions/deletions (indels) in any accession (Table 1). In total, 10,707,430 SNP-sites were available for analysis. When computing pairwise correlations of SNP-sites, we required the respective minor allele frequency to be greater than 5% at both sites. Allele frequency was computed relative to the number of accessions with unambiguous base calls (i.e. A, C, G, T only and with no ambiguous calls, such as N, allowed) at both sites, respectively. By requiring a minimum of 5% minor allele, we effectively selected older polymorphisms and excluded polymorphisms found exclusively in a small number of accessions (“private” polymorphisms). In total, approximately 5.7E+13 pairs of SNP-sites were inspected for evidence of correlated mutations.

**Table 1.**
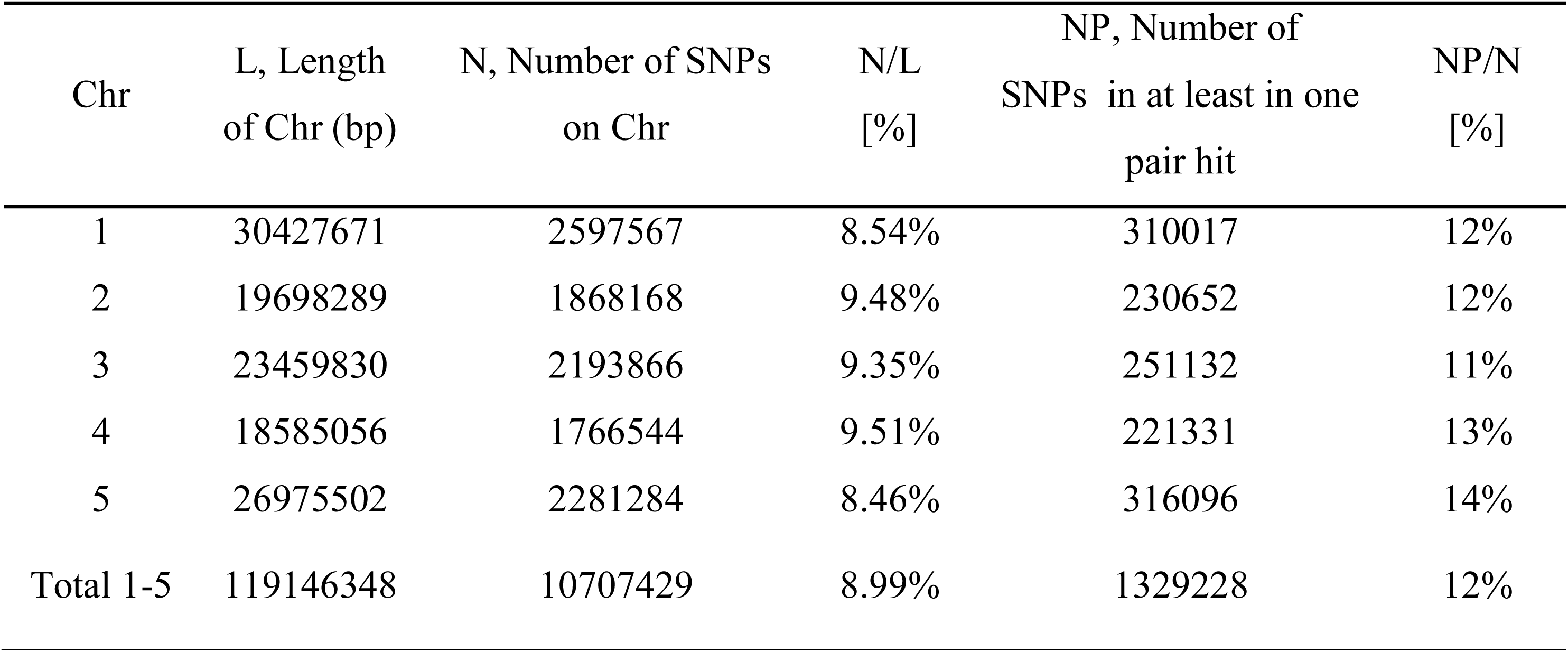
Summary of the numbers of SNPs per chromosome (Chr) and their relation to the size of chromosomes and the number of SNPs reported in pair hits.

### Adjusted Mutual Information (AMI)

To detect correlated mutations, the adjusted mutual information (AMI) was computed for every possible pair of polymorphic sites, using the analytical solution proposed by Romano et al., 2014 (18) (Eqs. 1–5). In essence, the AMI measures the agreement of the clustering of accessions based on alleles at two different sites corrected for their chance agreement. Given a specific polymorphic site, not all accessions in the dataset may show an unambiguous nucleotide call at this site, instead reporting Ns or other ambiguity codes. The lowest percentage of ambiguous base calls across all SNP-sites was determined as 4%; i.e. N-calls had to be taken into account for all pairwise comparisons. To detect covariance between two sites and without including ambiguous calls, the AMI was calculated for the intersection of accessions with definitive nucleotides called at both sites. We required this intersection to comprise at least 300 accessions. In total, approximately 5.7E+13 AMI-computations had to be performed, requiring efficient computing and program execution distributed across many CPUs. A C-program was implemented to perform the calculations in parallel on several cores per node using shared memory (C program available upon request). Note that the “adjustment” in the AMI-metric also corrects for differences in the lengths of base-vectors to be correlated and as caused by different occurrences of ambiguous calls (“N”s), thereby rendering pairs with different numbers of valid non-N accessions directly comparable.

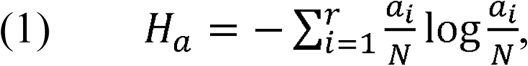

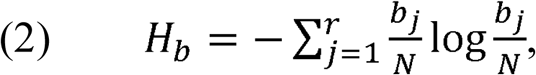

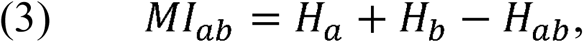

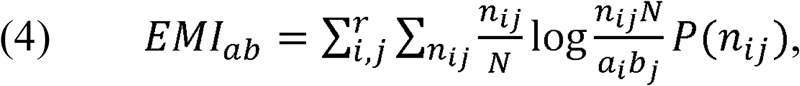

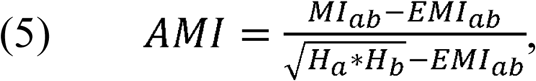
where *MI* is the Mutual Information, *EMI* the expected MI, *H_a/b_* is the entropy of variable *a/b*, respectively, and *H_ab_* is the joint entropy, *N* is the total length of the variable vector. In our case, *a_i_* and *b_j_* are the base counts of type *i=1..r and j=1..r* found at a particular SNP-sites *a* or *b*, respectively, *N* is the number of accessions with unambiguous base calls at both sites and *r* is the number of different alleles at a site (here, *r*=2, for biallelic SNP-sites), *n_ij_* is the number of accessions clustered together based alleles *i* and *j* at the two sites *a* and *b*. The probability *P(n_ij_)* is estimated from a hypergeometric distribution. AMI values range between 0, denoting random co-segregation of accessions, and 1, indicating perfect agreement of the allele-based partitioning at sites *a* and *b*.

Pairs of polymorphic sites that scored above an AMI value of 0.9 were considered correlated mutations (*pair hits*). The threshold value of 0.9 was chosen as it was found to be significantly larger than AMI-values that may arise by chance. This threshold was determined from an empirical permutation test, which was carried out using a subset of randomly selected 966,243 SNP-sites and vertically (across accessions) shuffled bases. The shuffling was repeated 1000 times. Then, computing all pairwise AMI values yielded a maximum AMI value of 0.88 with the bulk of AMI values at considerably smaller values (Supplementary Figure 1). Therefore, AMI >0.9 is a sensible threshold value. This sampling strategy to obtain a random reference distribution was chosen as the number of “N”-accessions is highly variable across different SNP-sites rendering analytical computations of the combinatory challenging.

With that many pairwise AMI computations for ~10Mill SNPs (5.7E+13), multiple testing appears to be a major concern at first. However, note that the probability of two SNP-sites randomly showing exactly correlated base changes is very small, and is, worst case 1.3E-25 for 300 non-N accessions and 5% (=15 accessions) minor allele frequency at both sites. In addition, with 1000 repeat runs of about ~1Mill shuffled SNP-sites, we performed about 10 times more random control simulations compared to actual comparisons. Thus, protecting against false correlations due to random chance can be safely controlled for. More critically, non-random effects leading to correlated mutations other than functional associations (e.g. via linkage or lineage) proved more challenging (see below).

### Annotation of pair hits

Pair hits (pairs of SNP-sites with AMI >0.9) were classified with regard to molecular and functional annotation based on the following criteria. First, for both of the SNP-sites forming a pair, the types of *genomic elements* to which they map were identified. Genomic element refers to the functional class of the encoded molecules (e.g. protein-coding or miRNA) at the respective sites or the type of local genomic region (e.g. intergenic, intron). A pair hit was then annotated as a particular *class* based on the combination of the genomic element annotations associated with the respective individual SNP-sites comprising the pair hit (*e.g*., genomic element (*A*): miRNA, genomic element (*B*): protein coding yields *class* (*A,B*): miRNA-protein coding). Class annotations with reversed order were combined into one class. Second, the sequence ontology classification associated with every SNP-site was mapped as well as the corresponding gene identifiers (AGI IDs) assigned if the site was found within known genes. This information was obtained from the VCF file from the 1001 genomes project (available http://1001genomes.org/data/GMI-MPI/releases/v3.1/1001genomes_snpeff_v3.1/1001genomes_snp-short-indel_only_ACGTN_v3.1.vcf.snpeff.gz). Pairs sharing identical gene IDs and genome element classification were considered intra-molecular/element pairs, otherwise, they were considered inter-molecular/element pairs.

### Accounting for linkage

SNP-sites at close genomic distances on the same chromosome will be linked because of the low probability of crossing-over (recombination) between the two sites and the associated reshuffling of alleles. Linked SNP-sites will yield to high AMI values, which most likely do not reflect functional associations, but a functional link is not strictly excluded either. Based on the observed distance intervals associated with pair hits (Figure 2), we set an effective distance cutoff of 10K bps to exclude linked sites.

### Sequence distance and average distance diameter: correction for lineage

The possibility that a pair of SNP-sites can have high AMI due to phylogenetic bias (lineage effect) was taken into account by the following approach. For all 1,135 Arabidopsis accessions, their respective pairwise genome-sequence-based distances were calculated based on 200,000 randomly selected polymorphic sites. The sequence distances were calculated using the software *Phylip* (19). For each pair hit, each of the two biallelic polymorphic sites split the set of accessions into two clusters, one with allele *a_i_*, the other with allele *a_j_*. For each accession cluster, the *Average Distance Diameter* (*ADD*) was calculated. ADD is the average pairwise sequence distance in the cluster of accessions sharing the same allele at the respective site (Supplementary Figure 2). ADD allows measuring how different the accessions in the cluster are with regard to their genomic sequence. In a given pair hit, the set of accessions may be divided into clusters because all accessions in each cluster originated from the same ancestor, and therefore, all its descendants have the same SNP mutation. Alternatively, the accessions in a cluster may be evolutionarily distant, but natural selection favors the specific allele observed at this site; i.e. positive selection preserved an allele in otherwise changing genomic sequences. When a cluster is explained by ancestry or phylogenetic relations, the ADD will be small because the sequences are related, and hence, similar. By contrast, when a cluster is explained by positive selective pressure, an allele is preserved even for distant accessions. In the latter case, this can happen possibly to compensate for a mutation in an interacting molecule, therefore the evolutionary sequence relations can be distant, and correspondingly, the ADD high. Each pair hit is composed of two SNP-sites, each SNP-site contains two clusters (*a_i_, a_j_*), each cluster has an ADD value that expresses the sequence divergence of that cluster. Thus, for every pair hit there are four ADD values, one for each accession cluster (two clusters) of each SNP-site (two SNP-sites). Given that the pair hits have high covariance (AMI >0.9), both SNP-sites involved in a pair hit cause the same (or almost the same) clustering of accessions. Therefore, the values of ADD differ between clusters of accessions, but not between SNP-sites in the pair hit (Supplementary Figure 2). The values for the ADD can vary depending on the sequence distance inside the clusters and several combinations of ADD between the clusters are possible. For example, cluster *a_i_* can have a small ADD, and cluster *a_j_* as well. This means that both clusters are likely caused by ancestry relationships. Cluster *a_i_* can have a small, and cluster *a_j_* a large ADD (or vice versa), then in cluster *a_i_* accessions are closely related, but in cluster *a_j_* they are not, in which case cluster *a_j_* is likely arose due to natural selection favoring that allele. Finally, both clusters can have large values of ADD showing a strong selective pressure for those SNPs in both clusters.

### Reducing redundant pair hits: correction for transitivity

AMI-values have been calculated for each pair of SNP-sites individually. As with any correlation measure, it may suffer from indirect correlations, also referred to as transitivity. When a SNP *A* is found to be correlated with both *B* and *C*, and based on a true mechanistic coupling, then *B* and *C* will appear correlated regardless whether or not they truly are associated mechanistically (Supplementary Figure 3). In our case, we corrected for transitivity based on profile vectors at SNP-sites; i.e. the pattern of base changes across the different accessions at a given site. For example, if detected that two molecules harbor SNP-sites with identical profile vectors, we may postulate a connection between them as they are correlated. If however, the very same pattern links these molecules to yet another molecule (or many others), we penalize this SNP-site according to the transitivity-value (or clustering coefficient) computed by the function “transitivity” of the R packages “igraph”. Thus, the transitivity value allows us to identify correlations that are less likely to originate from indirect correlations by enriching for sites with low transitivity value and reducing those that are redundant (high transitivity value). For isolated pairs of SNP-sites; i.e. both sites do not appear correlated to any other site, transitivity cannot be computed and an arbitrary transitivity value of -0.1 was assigned to both sites. Note that transitivity relates to unique patterns in SNP-sites; i.e. if molecule *X* is found correlated to molecule *Y* based two sites of identical SNP-patterns, and molecule *Y* is found correlated to molecule *Z*, but based on another SNP-site with a different pattern, we will not conclude *X* and *Z* to be linked as well.

### Testing for the effect of the introduced pair hit filtering steps using intergenic regions as reference

We tested for the effect of the introduced filtering steps (linkage, lineage, and transitivity) according to the following rationale. Given the set of actual SNP pair hits, we determined their associated characteristics with regard to annotation (pair hit classes as explained above, the genomic region/ molecule type). If the observed pair hits reflect true enrichment of particular classes, then a set of randomly obtained pair hits should show different characteristics.

The enrichment of particular classes was determined relative to pair hits that are unlikely to be functional. For the purpose of this study, pair hits in intergenic regions were assumed to be non-functional associations. Thus, the probability of finding pair hits in intergenic regions, with both SNP sites located in intergenic regions, was taken as the background probability.

The number of pair hits for a given interaction class was normalized by the number of possible comparisons amongst all SNP-sites in that class according to equations 6–10.

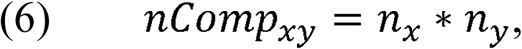

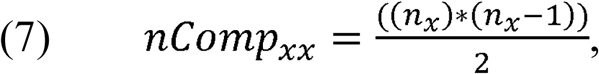

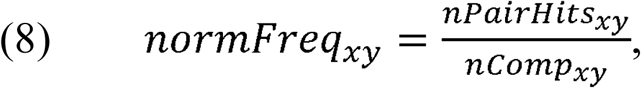

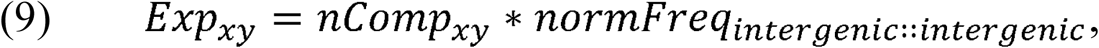

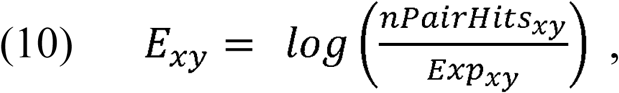
where *nComp_xy_* is the number of comparisons (SNP-SNP pairs) between two genomic elements *x* and *y*; *nComp_xx_*, the number of comparisons within the same genomic element *x*; *n_x/y_*, are the numbers of SNPs in genomic element *x* or *y*; *nPairHits_xy_* is the number of pair hits between genomic elements *x* and *y*, *normFreq_xy_* is the normalized count of the number of pair hits (*nPairhits_xy_*) over the number of comparisons (*nComp_xy_*); *Exp_xy_*, is the expected number of hits between class *x* and *y* using pairs between intergenic regions-intergenic regions as the reference. Enrichment (or depletion) factors (*E*) of pair hits relative to intergenic regions are expressed as logarithmic ratios (natural log) of observed vs. expected number of pair hits. In equations 8–10, if *x*=*y*, pair hits between the same type of genomic elements are captured, with equation 7 serving to compute the number of possible pairs.

For all possible combinations of genomic elements, the obtained enrichment factors, *E*, were compared to those obtained from random re-sampling by generating SNP pairs between two randomly selected SNP-sites. The random re-sampling aimed to destroy the covariance of the SNP-sites due to functional correlations, but to retain the characteristics of the observed pair hits that can be explained by confounding factors (linkage, lineage, and transitivity). If this random resampling produces similar enrichment profiles as observed for true pair hits with regard to their genomic class characteristics, the correlated mutations are unlikely to reflect functional associations, but merely are the consequence confounding factors. The random re-sampling was done by taking SNP-sites and repairing them randomly applying three conditions. (1) The distribution of the genomic distance between SNP-sites (number of bps between the SNP-sites) was forced to be identical for the actually observed pair hits and the re-sampled SNP-pairs. (2) The SNP-sites included in the resampling were filtered applying the same ADD (see above) as well as (3) the same cluster coefficient value (transitivity) thresholds. Thus, the filtering for confounding factors was kept, but the actual covariance was destroyed by randomly pairing up SNP-sites. In the re-sampling, only those SNP-sites, which were identified as part of pair hits, were considered, thereby eliminating SNP-sites from the original SNP-set that represent noise (regions that are difficult to sequence and, thus, rich in polymorphisms) or any other SNPs that are unlikely to represent any interaction. Furthermore, this selection renders the data manageable, as for ~5.7E+13 pairs, the fraction of N-base calls needs to be tracked resulting in prohibitively large data files. This approach of re-sampling only from those SNP-sites that were part of at least one pair hit was applied throughout this study whenever re-sampling was performed or the total number of comparisons was to be estimated.

The actual enrichment profile with regard to pair hit class was compared to the randomized data via the Pearson linear correlation coefficient, *r*. We applied this procedure for the various filtering steps with an effective filtering becoming apparent by a low correlation coefficient *r*. High correlations would suggest no difference of observed vs. randomized pair hit statistics, while low correlations would be indicative of differences, and therefore, reflect a signal in the observed data. In addition to the Pearson correlation coefficient, we computed correlation coefficients weighted by the number of pairs observed for every particular pair hit class by using the R package “wCorr”. Thereby, the influence of large, but not statistically supported enrichment factors associated with low numbers of occurrences (random fluctuations) is reduced.

### Annotation of known interactions, enrichment analysis, and semantic similarity of predicted protein-protein interactions

Three publicly available databases on molecular interactions were used to assign information to each pair hit, and to determine whether the polymorphic sites in a pair hit belong to a pair of molecules reported to interact. Protein-protein interactions (PPI) were identified using the AtPIN database (20), ANAP (21), and IntAct (22). In total, 18,288 unique proteins and their 187,313 interactions reported based on experimental evidence were used as a benchmark set after filtering the databases avoiding PPIs detected by predicting methods. Interactions between molecules not (yet) reported interacting, but with predicted interaction based on correlated mutations were tested for biological plausibility using the concept of semantic similarity. This method is based on gene ontology (GO) term annotation (23) and tests for an involvement of molecules (proteins) in the same biological processes.

The databases miRBase (24) and miRTarBase (25) were used to identify miRNA-mRNA-target interactions with 3,654 reported interactions serving as a benchmark test set.

To test for interactions at the chromatin level, we used the coordinates of chromatin loop formations at gene-resolution as reported in the study of Liu and collaborators in 2016 (26). SNP-sites located inside the coordinate intervals of DNA loops described to be in contact were considered as interacting polymorphic sites.

The enrichment of pair hits associated with interacting molecules (PPI, miRNA-mRNA, or loops) was tested by comparing the frequency of actual pair hits detected on molecules reported to interact to those obtained from random sampling. The random sampling was repeated 100 times and reported as a z-score statistic, with z-score computed as z-score=(f_a_-<f_r_>)/σ_r_, where f_a_ is the actually observed frequency, <f_r_> the mean random frequency, and σ_r_ the associated standard deviation. This comparison was done first using all the originally obtained pair hits against 100 samples of the same set, and secondly, using the filtered set (for lineage, linkage, and transitivity) compared also to 100 samples of this filtered set. As described above, in every random sample run, SNP-sites were randomly paired up creating the same number of pairs as in the original set and preserving the original chromosomal distance distribution, but reconnecting them randomly.

### Comparison with Hi-C data to detect long-range chromatin interactions

Long-range chromatin interactions have been studied in Arabidopsis using the Hi-C technology (27, 28). From Wang and collaborators, we received a table with the Hi-C data for the five Arabidopsis nuclear chromosomes containing Hi-C data presented as a 1,193 by 1,193 matrix of real-number values (28). This matrix consists of bins of genomic regions of size 100Kbp and using the TAIR9 (equivalent to TAIR10) genome assembly as reference. The Hi-C data were rendered binary by introducing a threshold of larger than 3 to indicate interactions, and otherwise no interaction, as recommended by the study authors (personal communication). To allow a comparison, we converted our pairwise and position-based data to a binned representation as well. The number of pair hits per cell (100Kx100K bp) was normalized by the number of possible comparisons between the two bins of length 100K bps, i.e. all pairwise comparisons of any SNP-site in each cell. A binomial test was implemented to define, which pairs of bins have more hits than expected. The expected probability of pair hits was estimated from the performed random sampling (see above). To account for linkage, only SNP-SNP pairs at distances greater than 10K or on different chromosomes were considered while also preserving the actual distance distribution of pair hits at larger distances. In addition, SNP-SNP pairs were filtered for lineage and transitivity (set Lk+Ln+Tr). Pairs of bins were considered significantly enriched for correlated mutations based on binomial testing with applied FDR correction for multiple testing (p_FDR_<0.05). Both data sets (Hi-C and bin-wise SNP correlation data) were tested for co-segregation using a Fisher exact test on the binary data; i.e. interacting yes/no, enriched for correlated mutations yes/no.

### Gene co-expression/ GO-term enrichment analysis

Gene expression information were obtained from NASCArray (29) with reported expression values for 20,807 unique nuclear encoded genes probed across 5,295 hybridizations covering a diverse set of conditions and using the ATH1 Affymetrix Arabidopsis gene-chip. Raw values were log-transformed and jointly normalized applying a quantile normalization using the “normalize.quantile” routine of the R package “preprocessCore”. Pairwise gene co-expression was assessed by computing Pearson correlation coefficients of the normalized expression values across all hybridizations.

Gene-ontology (GO) information was obtained from TAIR (30, 31). Enrichment of specific GO-slim terms in one particular gene set compared to another set was assessed by performing Fisher’s exact test of the respective term counts in the two sets to be compared. Associated p-values were tested for multiple GO-term testing based on the procedure introduced by Benjamini and Hochberg (32). Gene sets were not rendered unique; i.e. genes can be contained in the set engaging in positive and negative pairwise co-expression (with respectively differing gene partners) as well as, if a gene was reported in multiple pairings, its GO-term annotations were counted repeatedly effectively increasing their weights.

## Results

From an all-against-all comparison of all SNP-sites passing the selection criteria and across all considered 1,135 *Arabidopsis thaliana* accessions, 41,065,045 pairs of SNPs were detected at an Adjusted Mutual Information (AMI) level of above the employed threshold value of 0.9 and were considered significant correlations, henceforth referred to as *pair hits*. The total number of pair hits corresponds to 7*10^−5^% of all computed comparisons. Of all 10,707,430 polymorphic sites available for analysis, 1,329,228 (~12%) were part of at least one pair hit (Table 1). The distribution of individual SNP-sites in the five nuclear chromosomes of *A. thaliana* shows a specific segregation by genomic elements. Without correcting for gene-region size, many SNP-sites are found in protein-coding regions and transposable element genes. The SNP-density profile also reflects the chromosomal structure with high SNP densities near the centromeric regions, especially related to transposable element genes (Figure 1A). Most pair hits (84%) were found to be involved in inter-molecular interactions, with pair hits corresponding to protein-protein interactions being the most frequent correlated pair hit class (Figure 1B). However, upon normalization, we found that the most abundant pair hits relative to the corresponding number of comparisons were found on pairs of snoRNA-snoRNA, tRNA-tRNA, transposable element (TE)-TE, and TE-pseudogenes molecules (Figure 1C).

**Figure 1.**
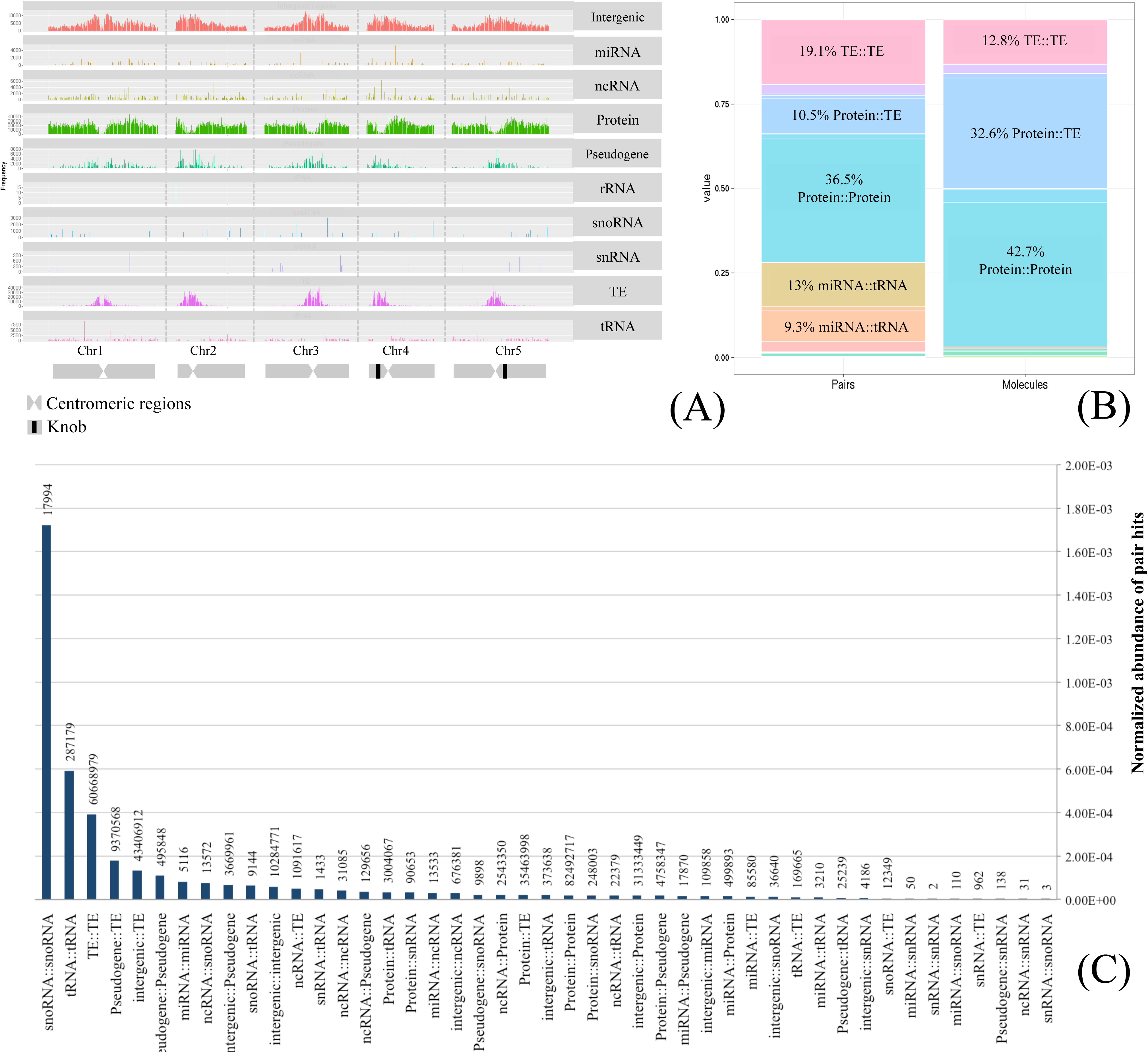
Summary of overall SNP and pair hits data and annotation characteristics. (A) Density of polymorphic sites along the genome as represented by the chromosome structure and molecule/genomic element type. (B) Percentage of pair hit annotation classes, counted as relative number of pair hits (left) and relative number of paired molecules (right). (C) Normalized abundance of pair hits representing the fraction of all significant SNP-SNP correlations carrying a particular type of class relative to all possible comparisons of any SNP-SNP site of this class. Numbers above the normalized frequency bars denote absolute counts of pair hits.

### Linkage disequilibrium

Correlated mutations will also reveal linkage effects, the joint inheritance of proximal SNP-sites due to reduced probabilities of crossing-over events, known also as linkage disequilibrium. Indeed, a pronounced increased frequency of pair hits at close genomic distances extending to a distance of up to 10K bps is evident (Figure 2). Most of the pair hits were found in close proximity along chromosomes with 47% of all pair hits (including inter-chromosomal pairs) detected within the genomic distance range of up to 10K bps on the same chromosome. Linkage disequilibrium appears lost at a distance of approximately 10K bps (Figure 2). This agrees well with previous reports on linkage effects in *A. thaliana* (33). Thus, linkage is an important source of SNP-SNP correlations. It does not require any functional and compensatory origin, and hence, needs to be accounted for in order to reduce false positive associations. Therefore, when comparing the observed correlation structure to random data, the random sampling was designed to reproduce the observed linkage decay (see Methods).

**Figure 2.**
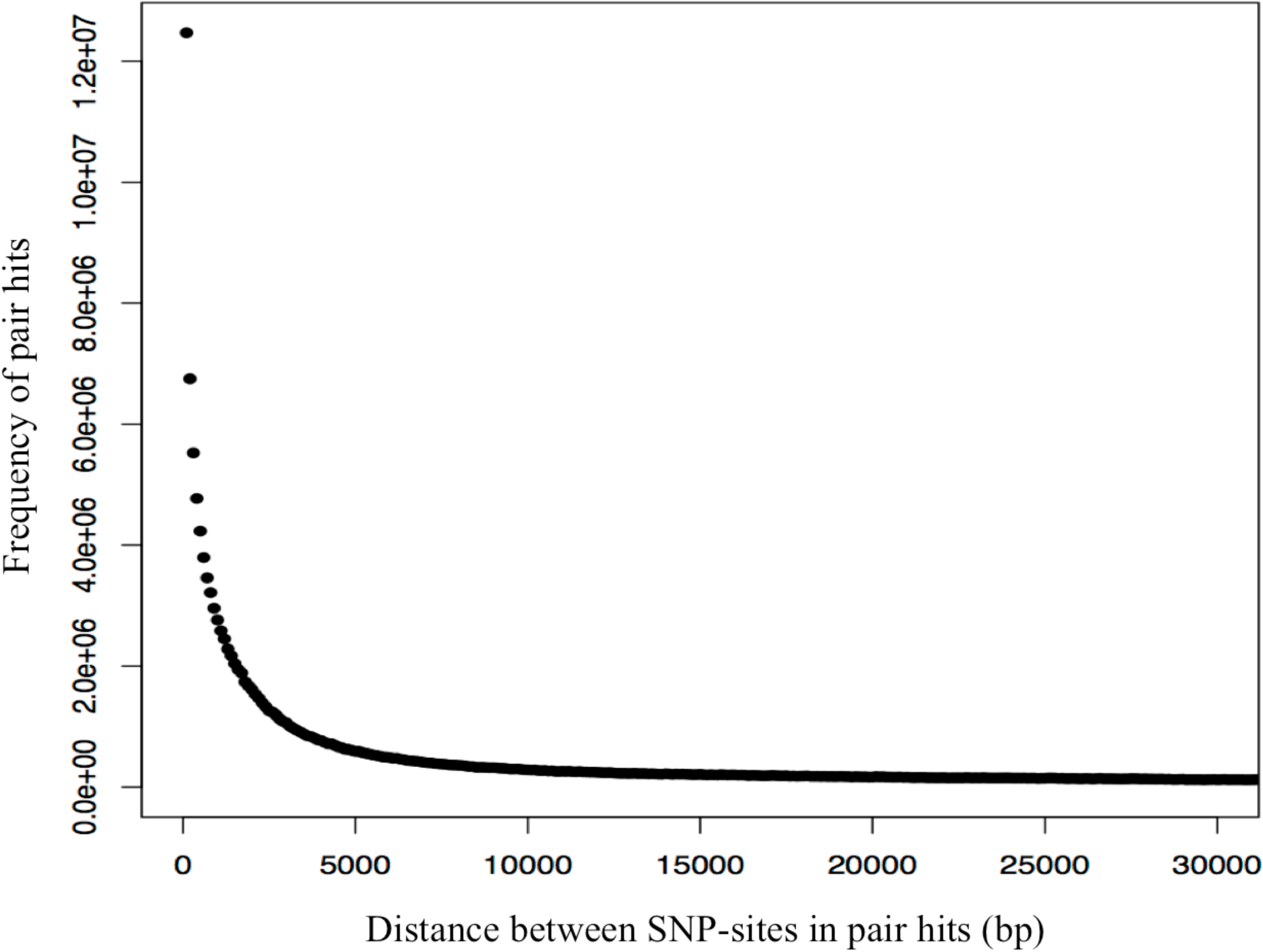
Frequency distribution of genomic distances in base-pairs (bp) of SNP-sites found correlated (pair hits) at bin width of 100 bps. To both, the numerator and the denominator, and arbitrary constant of 10 was added to avoid dividing by zero and to mitigate the noise due to small numbers.

### Correlation structure of the Arabidopsis nuclear genome

We inspected the global correlation structure of SNP-sites in the nuclear genome of *Arabidopsis thaliana* by comparing actual SNP-pair frequencies to two reference distributions. Most simplistically, the number of observed SNP-SNP pairs follows the density of SNPs alone. Evidently, and demonstrated already (Figure 2), linkage has a pronounced effect on the correlation structure and was accounted for by a distance-biased random re-sampling used as a second background distribution.

The heatmap of actual vs. expected pair hits indeed reveals regions of increased SNP-SNP correlations relative to both tested background frequencies (SNP-density alone, distance-biased to account for linkage) (Figure 3). Local clusters as well as long-range paired regions of elevated correlation can be identified. Local clusters primarily correspond to genomic regions around centromers. Long-range interactions also appear associated with these chromosomal segments.

**Figure 3.**
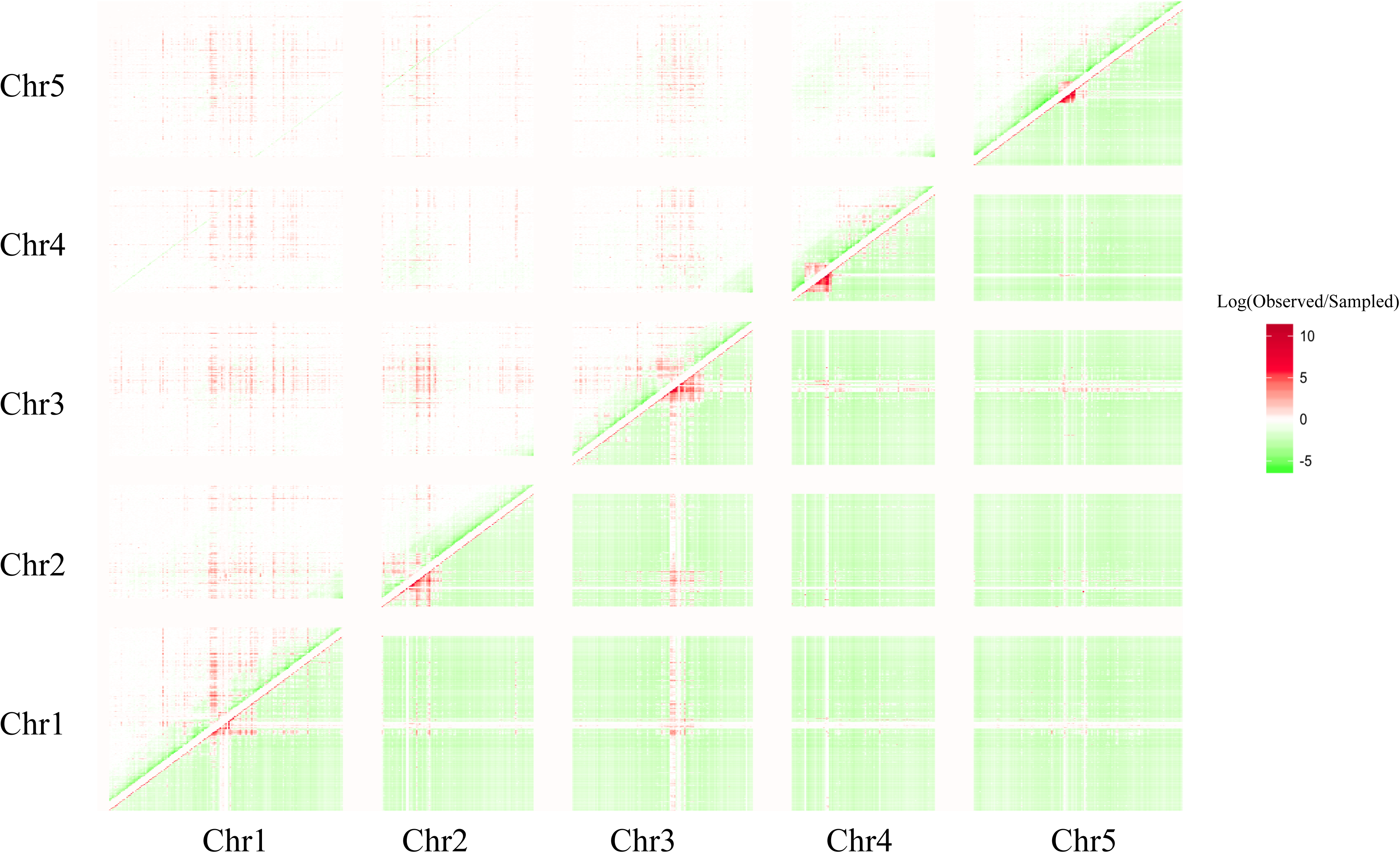
Heatmap of the natural logarithm of the ratio of the number of observed pair hits, *N_p_*, (correlated SNP pairs; counts per bins of 100K bps) and A) lower-right triangular matrix: the number of pairs estimated purely based on the density of SNPs, and B) upper-left triangular matrix: the number of random pairs forced to follow the same distance distribution (Figure 2). In A), the number, *N_p,r_*, of density-derived, random pairs per cell (*i*, *j*) (bin *i* (x-axis), bin *j* (y-axis) was obtained by *N_p,r_*=*Ni*Nj/T*, where *N_i, j_* are the number of SNPs in bin *i*, *j*, respectively, and *T* is a scaling factor to render to the total number of actual and density-based pairs identical. To both, the numerator and the denominator, an arbitrary constant of 10 was added to avoid dividing by zero and to mitigate the noise due to small numbers. Note that, also in the case of distance-biased re-sampling, the emerging clusters of elevated pair hit frequency do not reflect local SNP-density as the random re-sampling does consider local SNP density (see Methods).

When comparing pair hit counts to density-derived random pairings, again, the effect of linkage (Figure 2) becomes evident, as the diagonal (close proximity of SNPs) coincides with increased pair counts relative to random expectation at the expense of larger distances (lower than expected counts, Figure 3, lower-right triangular matrix). By contrast, the distance-biased resampling effectively accounts for the linkage effect (no elevated counts on the diagonal). Rather, except for the pericentromeric region, a relatively large (~2Mb) interval of seemingly lower than expected actual SNP-pairs is revealed (green band around diagonal in Figure 3, upper-left triangular matrix). This, however, may primarily result from the applied distance-bias-corrected re-sampling that effectively uniformly distributes the hugely increased pair frequency observed in the pericentromeric regions to all chromosomal regions (see Discussion on this point).

In addition to the position-local pair counts, both reference schemes reveal evidence of long-range and inter-chromosomal SNP-SNP correlations. In particular, the pericentromeric regions of all five chromosomes display pronounced increased pair frequencies over extended intervals and appear to be correlated across chromosomes. While within chromosomes, linkage may extend over longer regions due to decreased recombination rates near the centromere (16),across chromosomes, linkage cannot explain the observed high frequency of high correlations. Higher than expected counts of SNP-SNP pair correlations are also found outside pericentromeric regions.

Strikingly, vertical and horizontal stripes of elevated pair frequencies (red-colored cells in Figure 3, e.g. emanating from the interval 13–14Mb on chromosome 1, near the centromeric region, and extending across large genomic regions and even across different chromosomes suggest high frequencies of correlated mutations associated with selected regions of the nuclear genome. Upon further inspection, we concluded those correlations to result from the population structure present in the set of the 1,135 accessions. Correlated SNPs found in the regions associated with “stripes” were determined as those effectively segregating North American accessions from all others (European, Asian accessions). The colonization of the North American continent was reported to have been a relatively recent event (a few centuries ago) and associated with a fast geographic spread (34, 35). Thus, North American accessions are genetically more homogeneous and reveal a founder-effect, reported already in the seminal publication on the used set of 1,135 genomes to result in noticeable population structured evidenced by SNP-patterns (16). As a consequence, the particular alleles present in the founder population have been preserved and are revealed here as sites distinguishing them from other accessions leading also to their apparent correlation, which however, will likely have no functionally causal origin.

While this global picture does not provide any immediate functional insights, it does show that the actual correlation structure differs from random expectation and reveals the need to account for confounding factors, in particular, linkage and descent.

### Filtering out confounding factors

To identify pair hits that reflect functional associations via compensatory mechanisms, confounding factors, such as linkage (Lk), lineage (Ln), and transitivity (Tr), need to be taken into account. These biases have also been discussed in other studies (4, 8–10, 36). Hence, we aimed to determine their impact on our results and developed strategies to eliminate pair hits that are predominantly affected by linkage, lineage, and the transitivity effect. To detect whether the signal in the correlated mutations is originating from functional associations, we compared actual pair hits to carefully composed randomizations. The generated random re-parings of SNP-sites were designed to follow the same global linkage distribution and were subjected to the same lineage and transitivity thresholds as applied to the actual SNP-pairs (see Methods). For testing whether pair hits reflect functional associations as opposed to result from confounding factors, we pursued the following rationale: If observed pair hits are considerably different from random pair hits with respect to their genomic element type statistics (profiles as displayed in Figure 4), functional associations have likely been detected. By contrast, in cases of similar statistics of the genomic element types, the observed pair hits will predominantly be uninformative. Intergenic regions were assumed to be non-interacting and used as the background probability. For all pair hits, we found that interactions between RNAs (miRNA, tRNA, snoRNA, sRNA), transposable elements (TE), and pseudogenes have a probability of being correlated above what is expected based on the background probability, while many other classes occur less frequently (Figure 4A). Naively, we expected other types of interactions such as protein-protein interactions to be enriched relative to intergenic regions. Instead, they were found to be depleted. Furthermore, upon randomly re-sampling, a very similar enrichment profile was obtained as for the observed pair hits data (Pearson correlation coefficient, r=0.86, Figure 4A). Thus, when considering all pair hits, no strong evidence of functional relevance of correlated mutations was apparent.

**Figure 4.**
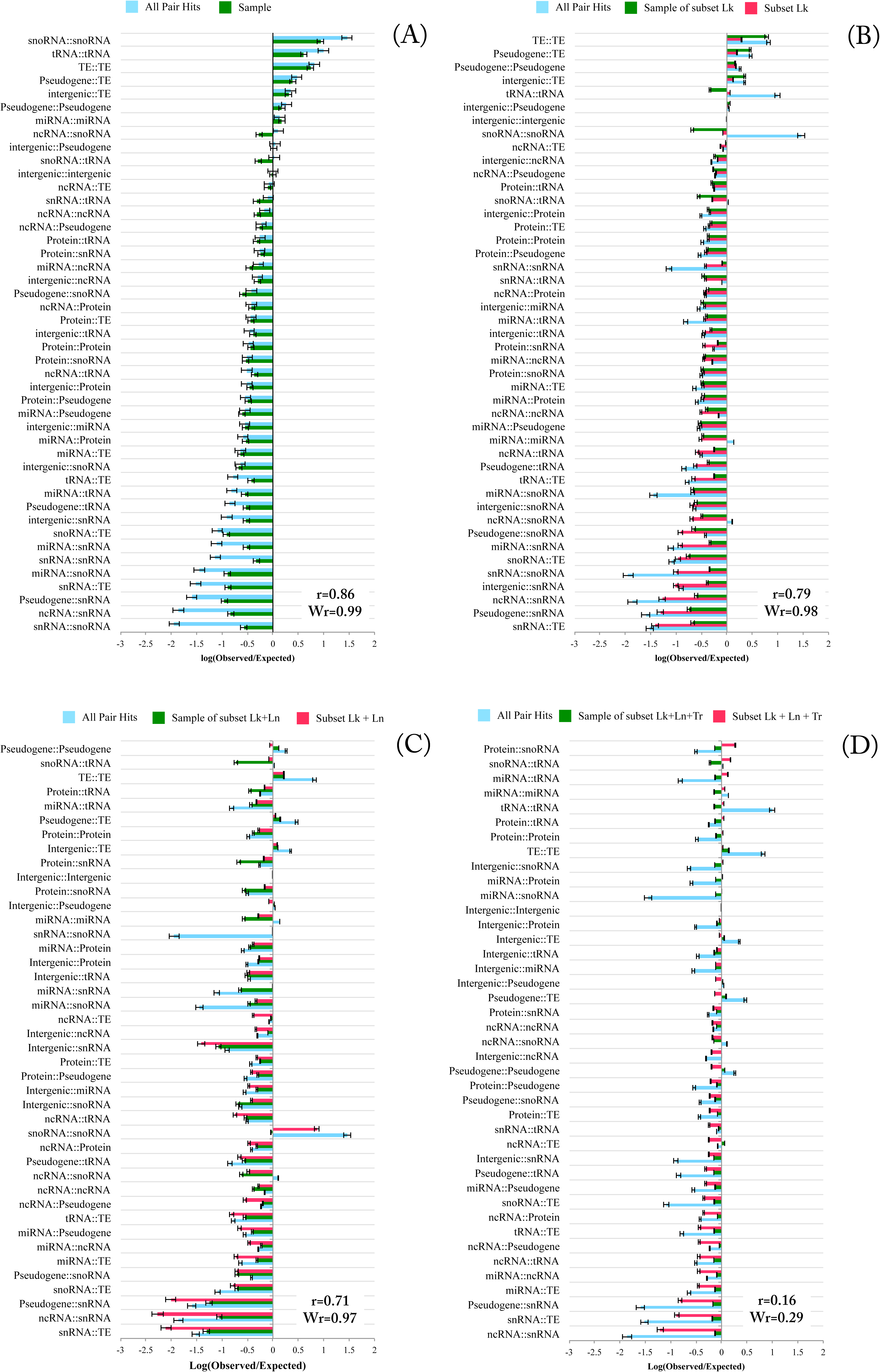
Enrichment (Eqs. 6–10, natural log) of pair hits in genomic classes using as reference intergenic regions-intergenic region correlations and compared to its resampling. (A) All pair hits. (B) *Subset Lk*, pair hits with more than 10K bps distance between them. (C) *Subset Lk* + *Ln*, pair hits with more than 10K bps distance between them and with associated values of average distance diameter (ADD) larger than the 75% percentile of ADD values. (D) *Subset Lk+Ln+T*r, pair hits with more than 10K distance between them, with values of ADD larger than 75% percentile of sequence distance, and with values of transitivity smaller than the 25% percentile of the cluster coefficient. Pearson correlation coefficients, r, of actual vs. random (sample) enrichment factors are given for each analysis. Wr-values correspond to correlation coefficients weighted according to the number of pairs entering the correlation. For comparison, raw enrichment values as reported in (A) are repeated in all graphs (light blue bars).

To account for linkage as a driver of correlated mutations, all pair hits with distances of smaller than 10K bps were discarded based on the observed distance dependence of pair hits (Figure 2). This subset is referred to as *subset Lk*. Again, enrichment of genomic classes represented in the set was tested relative to intergenic regions as the reference (Figure 4B). There was no pronounced difference in the enrichment factor distribution between the subset Lk and the complete results using intergenic regions-reference. Furthermore, when comparing the subset Lk to its random sampling, pair hits still exhibit a high correlation level (r=0.79).

Using different accessions from the same plant species can be expected to confer a strong phylogenetic bias to our results as the different accessions are closely related by descent. Therefore, it is essential to discriminate whether the detected covariance is functional or primarily reflects the phylogenetic history (lineage) and, thus, population structure. To account for phylogenetic bias, we measured all pairwise sequence distances between accessions and used the Average Distance Diameter (ADD) to assess the level of genome sequence diversity. The ADD was calculated among the groups of accessions that correspond to clusters of accessions based on the particular allele at a SNP-site (see Methods; for illustration, see Supplementary Figure 2). The rationale for this approach is to use SNP-sites resulting in clusters of accessions that correspond to high sequence diversity as this can be taken as evidence that natural selection may have favor a particular allele at a given site. Consequently, only pair hits with corresponding clusters of accessions displaying high diversity were considered. The top-75% percentile of the ADD diversity was used as a threshold. This subset was denoted as *subset Lk+Ln*, comprising 34% of the all pair hits.

The distribution genomic element enrichment of subset Lk+Ln displayed a different distribution compared to the pair hits obtained without any filters (Figure 4C). The distribution of subset Lk+Ln also showed increased deviation relative to its random sampling, albeit still at a high overall level, r=0.71. Thus, accounting for confounding factors still appears incomplete.

As the last explicitly considered confounding factor, we inspected possible effects from transitivity (Tr). Indirect covariances can result in overestimation of pair hits (see Methods, for illustration, see Supplementary Figure 3). Using pair hits with a smaller cluster coefficient, and hence, likely less affected by the transitivity effect, we aimed to reduce the number of redundant pairs. The 25% percentile of the lowest values of transitivity were considered for analysis (*subset Lk+Ln+Tr*, comprising pair hits with larger distance than 10K bps, the 75% highest ADD, and the 25% lowest transitivity values corresponding to 20% of the original set (8,549,704 pair hits)). The combination of the three filters yielded a distribution of the data that could not be reproduced by sampling. Sample and subset Lk+Ln+Tr have an associated correlation coefficient of r=0.16 (Figure 4D). Thus, following the rationale that actual profiles of genomic-class associations should not be reproducible by random sampling, we conclude that subset Lk+Ln+Tr may be enriched for functionally relevant correlations, and thus, may reflect molecular interactions. Interestingly, interactions such as tRNA-tRNA, protein-snoRNA, snoRNA-tRNA, miRNA-miRNA, miRNA-snoRNA, protein-tRNA, TE-TE, protein-protein (PPI), miRNA-protein, miRNA-miRNA, protein-tRNA, Protein-snoRNA were found overrepresented in this set relative to background (pairs of intergenic-intergenic regions). As these interaction classes appear biologically plausible, e.g. base-pair complementarity for RNA-mediated interactions or possible correlated amino acid changes in interacting proteins, or miRNA-target protein interactions reflecting the potential interaction of miRNA to the associated mRNA, the results obtained for subset Lk+Ln+Tr appear biologically reasonable. Likewise, pairings including pseudogenes were found at low levels, which can also be taken as evidence of the validity of our results as pseudogenes can be assumed non-functional. Note that the three introduced filtering steps were not applied sequentially, but were each based on the complete set and the intersection of those was taken. As the filtering reduced the number of pair hits, the correlation between actual and sampled pair annotations may decrease purely due to increased stochasticity (smaller numbers). However, the number of considered pair hits after filtering remained large. Furthermore, the enrichment factors obtained for the various classes appear biologically plausible. This view is supported further when inspecting protein-protein and miRNA-target gene interactions for the original and filtered set, where the filtered set yielded a larger fold increase of true interactions relative to random sampling (see below). To quantitatively account for the possible effect of small numbers, we also computed occurrence-weighted correlation coefficients (designated Wr in Figure 4). Observed similarly as for the Pearson correlation coefficients, a dissociation of class profiles of actual vs. random SNP-pairs is observed only when combining all three filtering steps (Lk+Ln+Tr, Wr=0.29), while the profiles remain closely correlated when considering all other filtering steps alone (Lk, Wr=0.98; Lk+Ln, Wr=0.97).

While the overall correlation structure after filtering (subset Lk+Ln+Tr) and as assessed by the heatmap of all-against-all chromosomal regions (Figure 5A, 100K bins) appear similar to the unfiltered pair hit set (Figure 3), pronounced differences can also be observed. Regions of increased density of correlated SNPs appear more condensed to large, but local clusters. The respective cross-peaks indicative of possible non-local interactions are relatively depleted except for correlations between chromosomes 2 and 3, for which the two respective local clusters also appear correlated among each other. Because of the introduced lineage and other filters, the “stripiness”, discussed above to result from the population structure and associated with North American accessions, has been reduced suggesting that non-informative correlations have been filtered out, albeit certainly not entirely.

**Figure 5.**
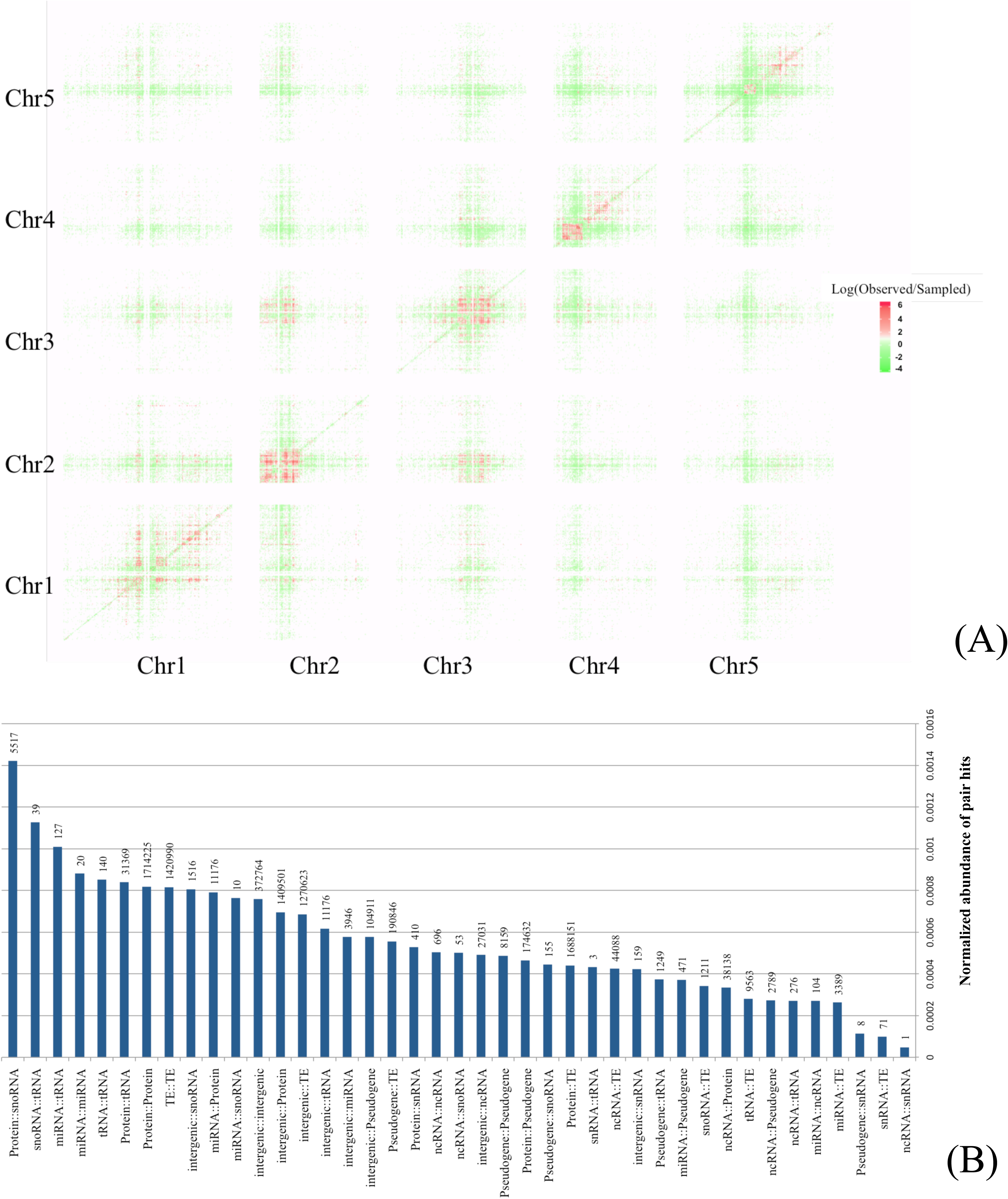
Summary of pair hit results obtained for subset Lk+Ln+Tr. (A) Heatmap of the natural logarithm of the ratio of number of observed pairs in the filtered data and the number of randomly sampled pairs, counts per bins of 100K bps. To both, the numerator and the denominator, an arbitrary constant of 10 was added to avoid dividing by zero and to mitigate the noise due to small numbers. (B) Normalized abundance of pair hits representing the fraction of all significant SNP-SNP correlations carrying a particular type of genomic element class relative to all possible comparisons of any SNP-SNP site of this class. Numbers above the normalized frequency bars denote absolute counts of pair hits.

Evidently, as the heatmap presents the averaged overview, individual pair hits can still be present across all chromosomes and distance intervals. Furthermore, inter-chromosomal regions corresponding to possible correlations between centromeric regions, observed with high SNP frequency (Figure 5A), appear depleted for pair hits after filtering, indicating a reduced probability of true functional interactions via compensatory mutations.

Pronounced differences are apparent with regard to relative abundance of pair hits for particular genomic element classes. As described before, classes that appear biologically plausible have moved to higher rank positions with regard to their enrichment factors (Eqs. 6–10) compared to the unfiltered set (Figure 4C compared to Figure 4A). For example, protein-protein interactions are ranked seventh after filtering, but were ranked 24th in the original set.

### Correlated mutations reflect known interactions

The subset Lk+Ln+Tr was tested for enrichment of known interactions between molecules. We focused on three types of interactions: protein-protein interactions (PPI), interactions between miRNAs and their target genes, and DNA-DNA interactions in chromatin loops and long-range chromatin interactions.

Concerning PPIs, 4,020 pair hits were found between proteins reported to interact; i.e. each of the two correlated SNP-sites were found present on proteins reported to interact. This set contained 290 unique gene-gene pairs corresponding to 642 unique protein-protein pairs as some genes encode more than one protein variant via splice variation. When performing 100 runs of random re-pairings of the same set of polymorphic sites (subset Lk+Ln+Tr), counts of pair hits on interacting proteins were found with a mean of 1,053+/-80 (s.d.). By comparison, actual pair hits were enriched 3.8 fold (z-score=36.9, p=1.9E-298) for known PPI relative to random pairings, and thus, may reflect functional associations between proteins. Without filters (all pair hits) the difference between the actual number of PPIs found with correlated mutations and the PPI found in the sampling was less pronounced and increased by a factor of 1.75 only (6,049,457 pair hits, z-score of 429.8, p=0). Thus, the introduced filtering steps proved effective in enriching for true interactions, but also significantly reduced the number of candidate associations, likely producing false negatives as well.

With regard to PPIs, the notion of compensatory mutations should have merit only in cases of nonsynonymous (i.e. amino acid changing) mutations. Synonymous (or silent) changes should not change the physics of interactions at all (but may affect other processes such as translation efficiency). Indeed, when examining SNP-SNP pairs on proteins known to interact, we found AMI-values associated with nonsynonymous-nonsynonymous changes to be significantly larger (mean=0.021, median=0.0096, Wilcoxon test p-value= 3.6E-118) than those obtained for synonymous-synonymous pairs (mean=0.018, median=0.007) (Figure 6). While significant, the magnitude of difference is small, possibly explained by synonymous changes being in turn correlated with nearby functionally relevant nonsynonymous changes or by providing a suitable genomic context for fixing nonsynonymous changes.

**Figure 6.**
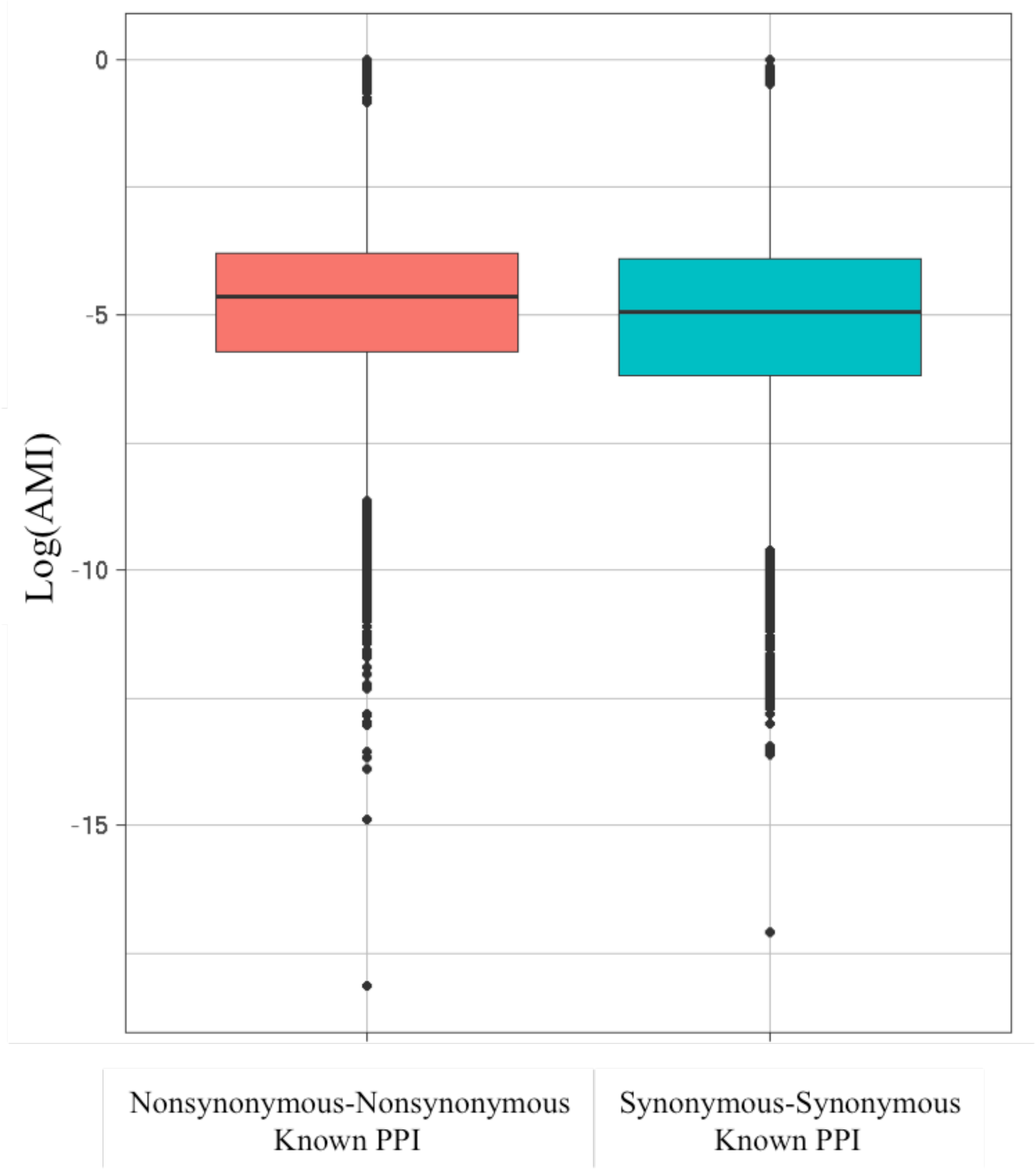
AMI-value distribution of SNP-SNP pairs on interacting proteins resulting either in nonsynonymous changes on both interacting proteins (mean=0.021, median=0.0096) or in synonymous changes (mean=0.018, median=0.0070), Wilcoxon-test p-value 3.63E-118, natural log.

Aside from protein-protein interactions, miRNA-protein-coding-mRNA are natural candidates molecular interaction types to harbor compensatory mutations, as for this type of molecular interaction, mutations directly translate into (almost) binary interaction decisions with base pairing possible or not. By contrast, mutations in proteins may either not result in any change of protein as they can be silent, and even if changing the amino acid, interactions between different amino acid residue types are not of binary character. Indeed, for miRNA-mRNA, we found 177 pair hits on 26 unique miRNA-mRNA molecule pairs corresponding to a 26.4 fold enrichment relative to random sampling (100 runs of random repairing, z-score=157.7, p=0). Again, when taking all pair hits; i.e. not including the filtering steps, the difference was less pronounced (fold=1.63, z-score=31.2, p=2.35E-214).

The statistic reported above was based on SNP-pairs (pair hits) and their distribution on interacting molecules. Next we examined the predictive value of pair hits with regard to molecule-based interaction predictions. While for both types of molecular interactions, the positive predictive value (PPV) was low (protein-protein interaction: 0.15%, miRNA-mRNA interactions: 0.7%; Table 2), for miRNA-mRNA pairs, the observed enrichment (odds reported by the Fisher exact 2×2 table test, i.e. counts of concordant/discordant pairs) was very pronounced (odds =5.36) and highly significant (p=2.5E-11). By contrast, for protein-protein interactions the enrichment of concordance was small (odds=1.16) and significance low (p=0.016).

**Table 2.** Predictive value of correlated mutations with regard to A) protein-protein and B) miRNA-mRNA interactions. Counts are based on a per-molecule basis, and were not based on SNP-pairs. Odds refers to the ratio of concordant (Yes/Yes, No/No) to discordant table entries (Yes/No, No/Yes).

| protein-protein |  | correlated SNP-pair |  |
| --- | --- | --- | --- |
|  |  | Yes | No |
| reported interacting | Yes | TP=290 | FN=50293 |
|  | No | FP=187023 | TN=37498722 |
| Odds (enrichment): 1.16 |  |  |  |
| Fisher exact test p-value: 0.016 |  |  |  |
| Positive Predictive Value (TP/(TP+FP)) = 0.15% |  |  |  |

B)
| miRNA-mRNA |  | correlated SNP-pair |  |
| --- | --- | --- | --- |
|  |  | Yes | No |
| reported interacting | Yes | TP=26 | FN=594 |
|  | No | FP=3628 | TN=444630 |
| Odds (enrichment): 5.36 |  |  |  |
| Fisher exact test p-value: 2.5E-11 |  |  |  |
| Positive Predictive Value (TP/(TP+FP)) = 0.7% |  |  |  |

### Gene-loops and chromatin interactions

Lastly, we probed for correlated mutations to inform on physical interactions at the DNA/chromosomal level testing both short-range gene loop formations (26) as well as long-range chromatin interactions (27, 28) reported in Arabidopsis and based on Hi-C data.

In reported DNA-DNA contacts in gene loops, we found 60 pair hits (fold=2.8 relative to the mean obtained from 100 runs of random re-pairings, z-score=8.74, p=2.14E-18). In this test, the introduced distance filter of 10K bps eliminated most of the reported gene loop contacts. When including all pair hits (without any filtering), we detected 432,870 pair hits, 4.3 times more than the mean obtained from the 100 runs of random re-pairings of the same number of original pair hits (z-score=1,102, p=0). Note that, while there will be substantial linkage within 10K bps, and thus many pair hits (Figure 2), in the random repairing the distance-dependent frequency distribution was preserved to account for linkage. Thus, correlated mutations also reflect on chromosomal gene loop contacts.

Next, we tested whether correlated mutations reflect long-range chromatin interactions. Specifically, we asked whether sites of physical contacts between distant chromosomal regions overlap with pairs of genomic regions rich in correlated mutations. Both, Hi-C data informing on physical chromosomal contacts and pair hits reflecting correlated mutations were binned into intervals of 100K bps, making each pair (cell) of 100K-100K bins the basic interacting unit. We found a significant overlap of long-range chromatin interaction cells detected using Hi-C technology and cells with significant numbers of pair hits (Fisher’s exact test, *p-value* < 2.2e-16) (Figure 8A). The correspondence appears particularly strong near pericentromeric regions, both within and across different chromosomes. In comparison to the Hi-C data, a larger number of cells appear to be interacting in our pair hits data (blue cells in Figure 8A) as our data can also reflect different levels of interactions, not only at the chromosomal level, and/or because of differences in thresholding. However, the significance of the Fisher test suggests that correlated mutations may indeed reflect long-range chromatin interactions.

In the pairwise regions with concordant evidence of interaction revealed by both SNP-SNP correlation (pair hits) and Hi-C data, 29.9% of the genomic classes are TE-TE genes pair hits, 24.9% intergenic regions-TE and 20.6% Protein-TE (Figure 8B), all found at higher frequencies than in regions not showing agreement of pair hit and Hi-C data (discordant cells, Figure 8B, TE-TE 19%, p<2.2E-16; intergenic-TE 12.%, p<2.2E-16; Protein-TE 10.5%, p<2.2E-16 with p-values obtained from Fisher’s exact test compared to concordant regions). Moreover, there is an enrichment of intergenic-intergenic pairs in chromatin interacting regions concordant with our data compared to discordant cells (4.8% compared to 1.6%, p<2.2E-16). By contrast, pairs of the class protein-protein interactions are depleted, explained by the paucity of protein coding genes in pericentromeric regions (Figure 1).

### Prediction of novel protein-protein interactions

As pair hits were indeed found to reflect molecular interactions between proteins, it can be assumed that also those pair hits that link two different proteins that are not reported to interact based on the currently available information may, in fact, interact. To test for this possibility and taken as an indirect evidence, we checked whether two proteins connected via at least one identified pair hit, but not reported to interact, are involved in the same biological process as captured by their GO-term annotation. Indeed, the semantic similarity of the putative 471,002 PPIs based on pair hits proved significantly higher semantic than obtained for a random sampling of protein pairs (Figure 7, Wilcoxon test p-value < 2.2e-16).

**Figure 7.**
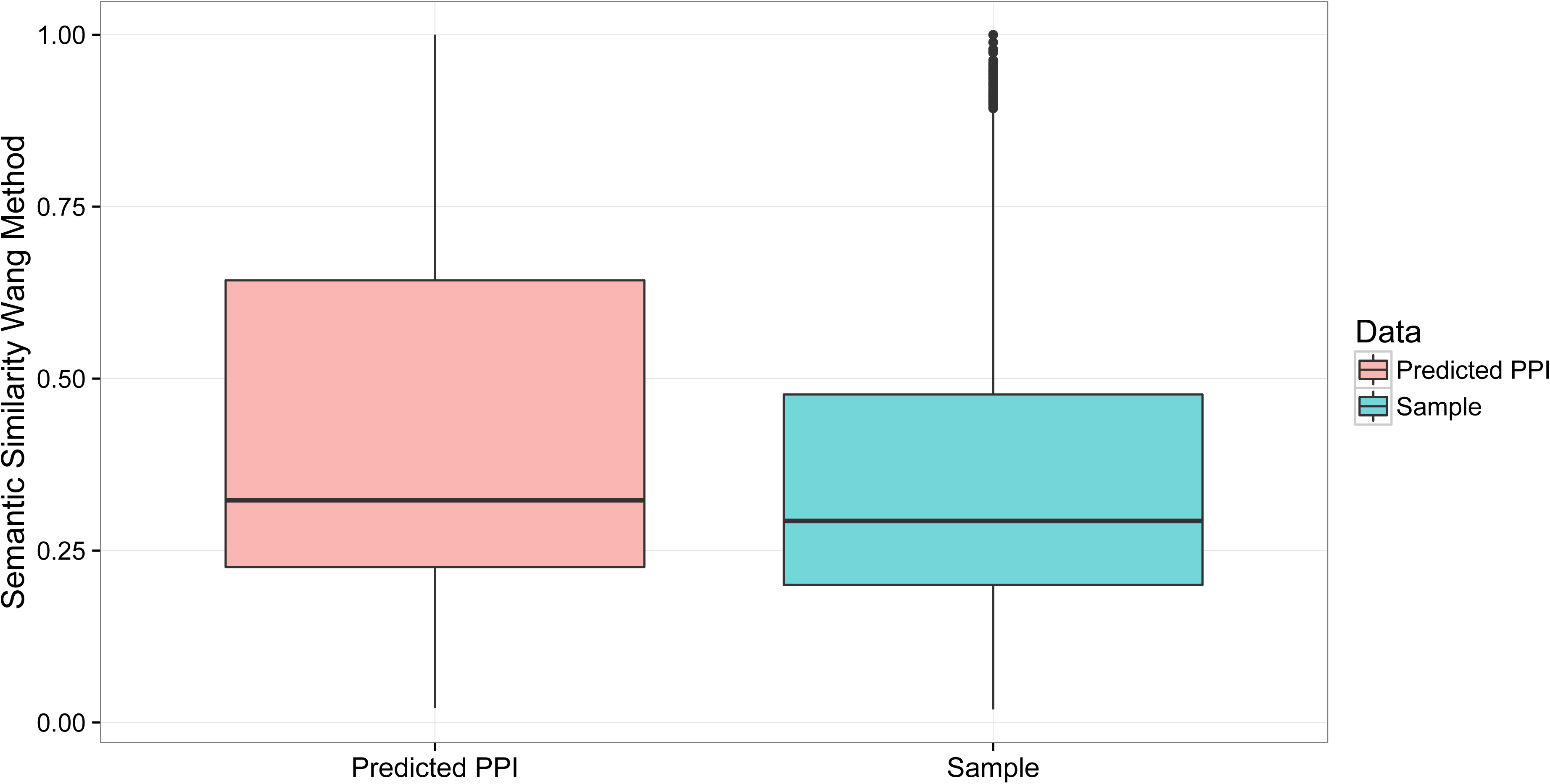
Boxplot of the semantic similarity of proteins predicted to interact based on a pair hit linking them compared to a set of protein pairs formed randomly. In this statistic, only those protein pairs were considered that are not reported to interact using the available PPI information (Wilcoxon test p-value < 2.2e-16).

In addition to novel protein-protein interactions, Table 3 lists the pair counts of all other molecule types predicted to interact based on detected pair hits (filtered set) with one and, imposing a stricter filtering, five or more showing them as correlated. Actual molecule pairs associated with the latter are available as Supplementary File 1. Of note are the highlighted predictions for the types detected to be enriched relative to expectation (Figure 4D).

**Table 3.**
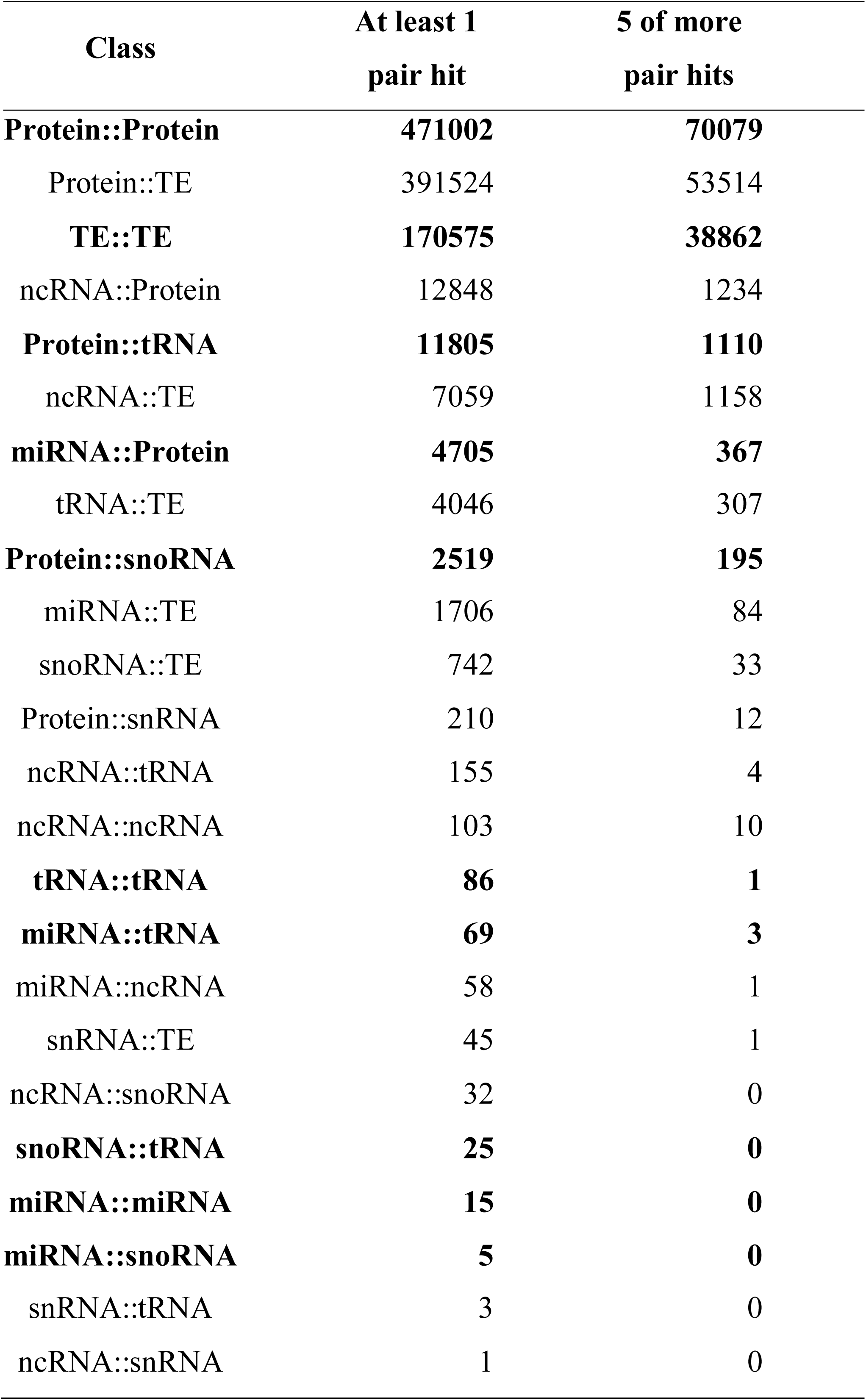
Predicted interactions of molecules based on the presence of at least one/ five pair hit(s) revealing them as correlated. Highlighted bold are the molecule type pairs identified as enriched for pair hits in the genome-wide screen and after Filtering (Figure 4D) (Transfer RNA (tRNAs), small nuclear RNA (snRNA), Small nucleolar RNAs (snoRNAs), non-coding RNA (ncRNA), microRNA (miRNA), protein coding sequences, Transposable Elements (TE)). Actual molecule pairs predicted to interact based on five or more pair hits are available as Supplementary File 1.

| Class | At least 1<br>pair hit | 5 of more<br>pair hits |
| --- | --- | --- |
| **Protein::Protein** | **471002** | **70079** |
| Protein::TE | 391524 | 53514 |
| **TE::TE** | **170575** | **38862** |
| ncRNA::Protein | 12848 | 1234 |
| **Protein::tRNA** | **11805** | **1110** |
| ncRNA::TE | 7059 | 1158 |
| **miRNA::Protein** | **4705** | **367** |
| tRNA::TE | 4046 | 307 |
| **Protein::snoRNA** | **2519** | **195** |
| miRNA::TE | 1706 | 84 |
| snoRNA::TE | 742 | 33 |
| Protein::snRNA | 210 | 12 |
| ncRNA::tRNA | 155 | 4 |
| ncRNA::ncRNA | 103 | 10 |
| **tRNA::tRNA** | **86** | **1** |
| **miRNA::tRNA** | **69** | **3** |
| miRNA::ncRNA | 58 | 1 |
| snRNA::TE | 45 | 1 |
| ncRNA::snoRNA | 32 | 0 |
| **snoRNA::tRNA** | **25** | **0** |
| **miRNA::miRNA** | **15** | **0** |
| **miRNA::snoRNA** | **5** | **0** |
| snRNA::tRNA | 3 | 0 |
| ncRNA::snRNA | 1 | 0 |

### Co-expression of gene pairs linked via correlated mutations

In addition to direct physical interactions, functional links between two genes linked via correlated mutations could also become evident as correlated gene expression behavior. Exploiting a large compendium of gene-expression microarray data (see Methods), we indeed detected differences of the levels of gene-co-expression for gene pairs linked pair hits compared to gene pairs from a random pairing of SNPs (Figure 9). Here, and to allow for the effect of possible regulatory effects, we considered all SNP-locations within gene; i.e. including also non-CDSs sites such as introns or UTR. However, contrary to expectation, we did not find correlation levels to be shifted to larger positive values, but instead, a shift to larger absolute values. This result was even more pronounced when selecting only those gene pairs with five or more pair hits linking them (medians of absolute pairwise gene expression Pearson correlation coefficients of random gene pairs=0.086, median of gene pairs linked by at least one/five pair hit=0.091/0.097, p=7.7E-87/2.46E-169 (Wilcoxon test)), albeit the magnitude of difference was small. Thus, the expression of genes linked via pair hits is notably different from their random expectation.

To understand the unexpected increase of not only positive, but also negative correlation levels between genes with correlated mutations, we performed a GO-term (functional terms) enrichment analysis of gene sets with genes engaging in pairwise positive correlation (*r* > 0.2) and contrasted those to a set of genes with pairwise negative co-expression regulation (*r* < −0.2). In line with expectations, genes engaging in positive pairwise gene co-expression and with at least 5 or more pair hits were enriched for the GO term “protein binding”, p_FDR_=1.8E-11 (other GO terms with p_FDR_<0.05: “receptor binding or activity” p_FDR_=5.8E-4; “nucleotide binding”, p_FDR_<2.6E-3). Thus, the notion of correlated mutations reflecting functional, and primarily physical, interactions seems validated as positive gene expression correlation is immediately consistent with this mode of interaction. By contrast, among genes with negative expression correlations, but also linked via at least 5 pair hits, no single GO term reached the same level of significance as observed for “protein binding” in the positive correlation set. Instead, GO terms “DNA or RNA binding”, p_FDR_=1.97E-3; “unknown molecular function”, p_FDR_=1.97E-3; “transferase activity”, p_FDR_=2.6E-3; “transporter activity”, p_FDR_=3.1E-3; “nucleic acid binding”, p_FDR_=1.2E-2; “other binding”, p_FDR_=1.2E-2, were obtained at borderline significance levels.

## Discussion

The concept of detecting and exploiting correlated mutations as indications of physical or functional interactions has been appealing from its inception (37–39). Its underlying rationale appears straightforward and convincing. Furthermore, the basis for its successful application - the availability of broad sequencing information - is growing rapidly. The power of this approach has also been demonstrated by its successful application in predicting structural aspects or proteins (e.g. (8, 11)) and RNA (36). At its core, the method requires evolution to have acted to select for compensatory mutations. Thus, sufficient evolutionary time needs to have elapsed in order for evolution to take effect. Therefore, previous studies on correlated mutation aimed to increase evolutionary distances between the considered genomes and included different species only.

Here, we set out to test and apply the concept of correlated mutations within the same species (*Arabidopsis thaliana*). Evidently, the considered genomes are more closely related with less evolutionary time separating the different Arabidopsis accessions as compared to different species, and therefore, the selective forces of evolution will have had less time to act. It has been estimated that *Arabidopsis thaliana* diverged from its ancestors around 5 MYA (40); i.e. relatively recently compared to the ~450 million years of land plant evolution (41). North American accessions are supposed to have colonized their current habitats as recently as a few centuries ago (34, 35). Thus, it was not obvious whether the concept of correlated mutations can be used successfully when applying it to within-species variation. Nonetheless, by implementing a series of filtering steps and by performing appropriate controls, our study suggests that correlated mutations can also reveal physical, or broadly functional, interactions between molecules or genomic regions even within a species. Correlated mutations were found to be enriched in molecule pairs known to interact. Thus, this confirmation also implies that this approach may also have predictive value, even though the number of false positive predictions can be expected to be high (Table 2). Our study also reveals the overall genome-wide correlation structure of the Arabidopsis genome and required implementing an efficient computation scheme rending the computation of around 10^13^ comparisons possible. Subsequently, we shall discuss methodological aspects as well as the biological implications of our results.

Mutual information - the metric employed in this study – is a mathematically robust method to detect covariance between categorical variables (here, alleles). However, its power to detect functional associations has also been questioned due to different confounding factors (4, 8, 10). Two positions in an alignment can be covarying because they descended from the same ancestor in which two random mutations appeared (lineage) or because they are inherited jointly due to proximity on the chromosome (linkage) (33). Indeed, both effects became evident in this study as well (Figure 3). In addition, methods for measuring covariance frequently fail to properly account for the transitivity effect caused by indirect correlatoins, thereby overestimating the number of interactive pairs (8, 10). Additionally, sequencing errors could result in false and repetitive SNPs. This could lead to a substantial overestimation of the number of pair hits due to the transitivity effect, because all repetitions appear to be covarying with each other. To address these difficulties, we devised as series of filters. By maximizing genomic sequence distance within the splits of accessions introduced by the two alleles at a given site, we aimed to reduce the lineage effect. By penalizing sites with a high clustering coefficient, we diminished the influence of the transitivity effect. Linkage was avoided by only considering pairs at a distance of 10K bps or larger. As a consequence, the annotation characteristics of the actually detected pair hits differed significantly from its random control (Figure 4D), which can be taken as evidence that the filters have yielded non-random SNP-correlations. As furthermore, the fold enrichment of known interactions detected by pair hits increased after filtering, we can also conclude that the effect of the introduced filtering steps was not only leading to non-random pair hits, but that those are enriched for true associations. We also tested for the possibility that sequence repeats may result in increased pair hit frequencies due to sequence read mapping mistakes. However, no such correlation was evident (using repeats of length 15bps spaced at 500 bps and testing exemplary chromosome 1; not shown.)

Our reasoning to conclude that we detected functional associations relied on comparisons to random controls. When generating random background pair distributions, we also aimed to avoid spurious associations by accounting for confounding factors. First, by maintaining the distance distribution between pairs in the random sampling, we tested if the enrichment was related to linkage, recombination patterns, gene conversion, chromosome structure or another factor related with the location of polymorphisms that could be influencing our findings. Without any filtering, these location-associated effects were indeed found to explain to large extent the observed correlation structure (Figure 4A), and thus needs to be taken into account. However, randomly sampled pairs with low transitivity and high sequence diversity in combination with maintaining the distance distribution showed little correlation with the real, filtered results (Figure 4D). This suggests that the enrichment observed after filtering is neither related to the distribution of the polymorphic sites in the genome, nor to the influence of phylogenetic biases, nor to the transitivity originating from indirect covariation. Consequently, we feel supported in the conclusion that the observed covariance will reflect functional association. The recognition of linkage, lineage, and transitivity as strongly influencing factors is also important for the prediction of new interactions. However, these factors do not preclude true functional association, and thus, by filtering for them, truly functionally correlated sites may have been lost upon filtering.

To account for linkage, we generated random SNP-pairs to follow one distance distribution, expected to be globally valid, and introduced a distance cutoff of 10K bps. However, near the centromers, linkage has been reported to show increased linkage (16). Consequently, a separate treatment of pericentromeric regions with regard to linkage seems indicated. While treating the background distribution uniform across all chromosomes and regions did initially, i.e. before applying the filters, indeed result in noticeable biases (green band along the diagonal in Figure 3, upper triangle), the effect was largely eliminated when applying all filters (Figure 5A). It needs to also be noted that given the SNP-data, it is not possible to clearly separate out linkage-effects from possible true functional correlations, for which, indeed, we found evidence (chromosomal contacts, Figure 8).

**Figure 8.**
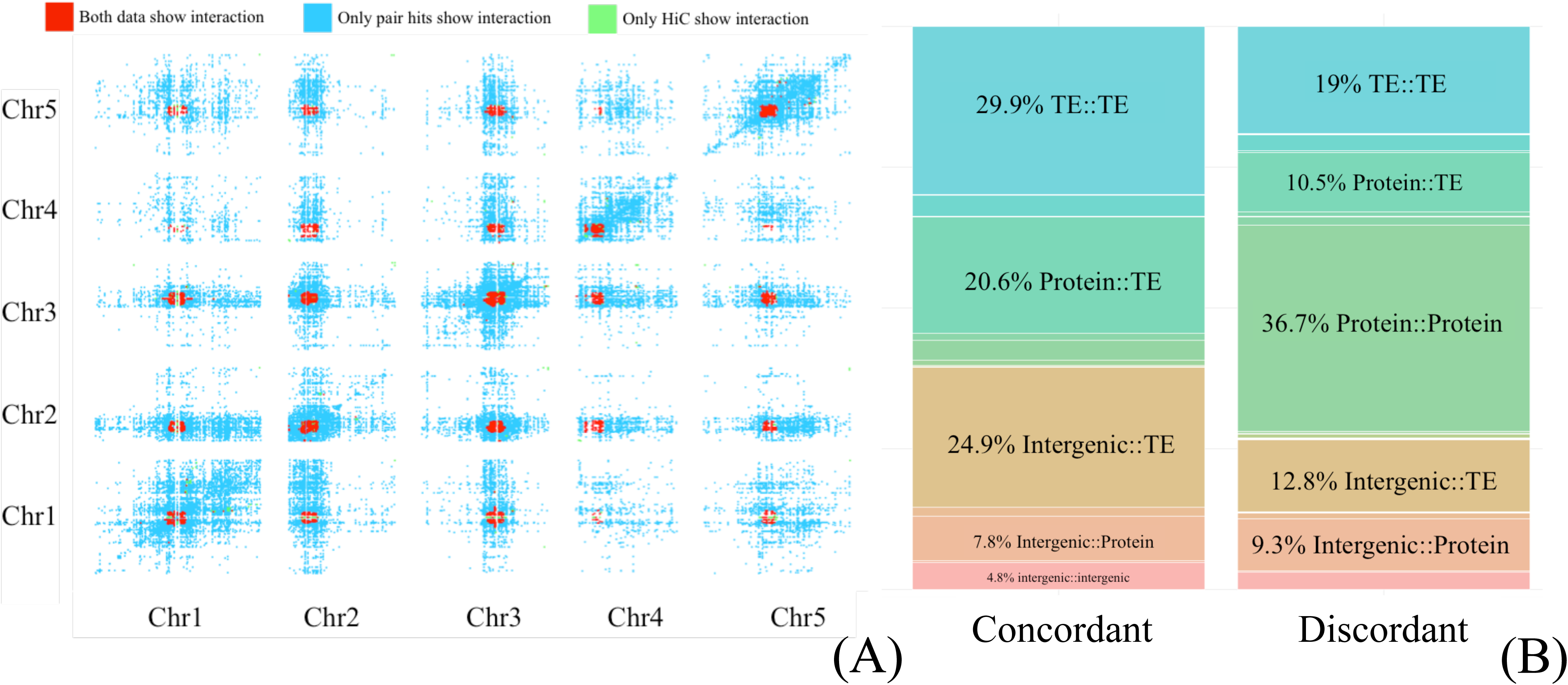
(A) Heatmap of genomic regions considered interacting. Cells (pairwise bins of 100K bps) that have significantly more pair hits than randomly expected are considered correlated and colored blue. Cells (same bin size of 100K bps) considered physically interacting based on Hi-C data shown in green. Overlapping cells for both sets are colored red. (B) Percentage of pair hit annotation classes, counted in paired regions that appear to be chromatin interactions as well as enriched for correlated mutations (left panel, “concordant”) and counted in regions that are not chromatin interactions (right, “discordant”).

**Figure 9.**
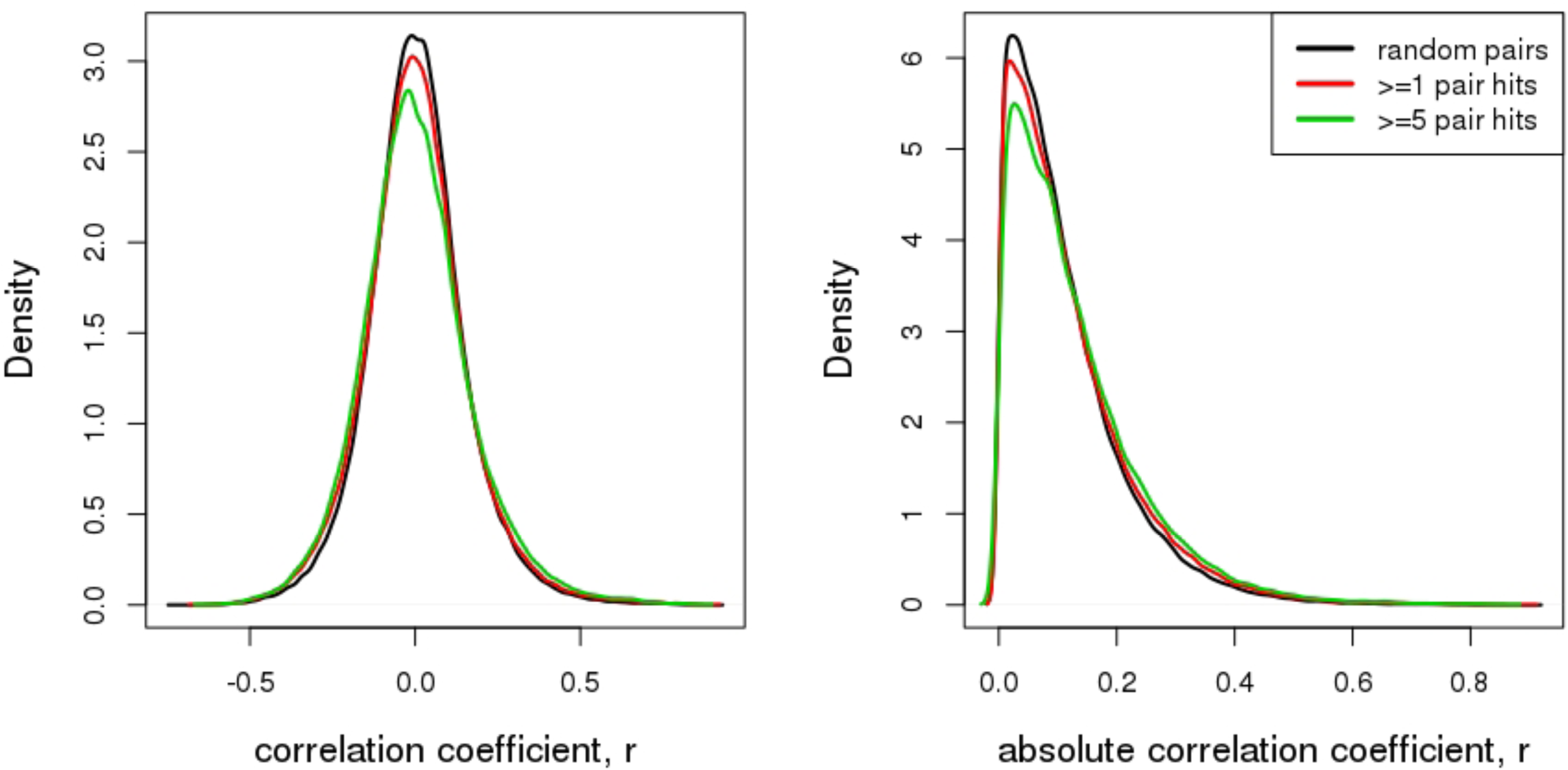
Pairwise gene expression correlation of transcripts encoded by gene pairs linked via correlated mutations (pair hits) and as measured by raw (left) and absolute (right graph) Pearson correlation coefficient, r. Considered gene-pair sets were linked by at least one (294, 479 gene pairs with available gene expression information for both genes) or five pair hits (56, 718 pairs) and compared to a set of randomized pairings (172, 443 gene pairs) of the same set of SNP-sites. All SNP-locations on genes were considered; i.e. also including non-CDS regions such as introns or UTR.

Natural selection that favors specific adaptations to specific environmental conditions represents another origin for covariance, also referred to as purifying selection (42, 43). In essence, environment acts as an independent factor producing indirect correlations between SNP-sites that are themselves not directly functionally linked. For example, accessions in hot climates may show specific mutations increasing their fitness relative to those in colder habitats. Those sites would be detected as correlated, but they are caused by a third factor, the environment. Furthermore, the environment may not only introduce independent changes, but may also affect the fixation of compensatory mutations. In a yeast model of a particular human disease, it was demonstrated that environmental conditions influence the fixation of compensatory mutations (44). However, it has been reported that purifying selection in coding-regions of plant genomes is virtually absent (45). This would suggest that the extent of correlation due to environmental factors and purifying selection may be small in Arabidopsis. However, it needs to be cautioned that seemingly neutral mutations may reflect unrecognized adaptations. Furthermore, it has been found that at least 5% of noncoding regions in *A. thaliana* are under purifying selection in the family of Brassicaceae (46). And although in the original publication reporting the 1,135 genomes used here, evidence of environmentally driven selection of alleles was demonstrated for a small set of SNPs (16), the authors cautioned that any attempt to infer selection of particular alleles is inevitably obscured by population structure effects. Therefore, indirectly correlated mutations induced by external factors cannot be ruled out to contribute to the set of pair hits reported in this study.

When using closely related genomes as done in this study, detecting correlated mutations necessarily becomes challenging for reasons of direct ancestry and linkage. However, it also bears the advantage of allowing direct vertical alignment comparisons across the genomes without necessitating establishing homology between different molecules in different species. Yet, large deletions/ insertions and rearrangements may happen also within species and may constitute a source of error of our study (47, 48).

Epistasis plays an important role in the fixation of compensatory mutations and is pivotal for molecular evolution (44, 49). This phenomenon allows the fixation of deleterious mutations if compensated by other mutations, or generally by the genetic background of the organism. Therefore, in essence, our effort to identify correlated mutations can also be seen as yielding epistatic events and reveals the extent of this mode of evolution. Epistasis may also explain the unexpectedly small difference between the AMI of nonsynonymous-nonsynonymous and synonymous-synonymous pair hits in known PPI (Figure 6). While not changing the amino acid, synonymous changes may provide the conducive background for a nonsynonymous change via an as yet unidentified mechanism.

Aside from addressing the question as to whether even within-species variation reveals evidence of compensatory mutations, ideally, correlated mutations can be exploited for predictions. While for miRNA-mRNA interactions, statistics derived from both, SNP-pair-based as well as molecule-based, resulted in significant positive predictive values, protein-protein interactions were essentially evident only when considering SNP-pair statistics, but not at the level of molecule-molecule interactions (Table 2). Thus, on protein-protein pairs reported to interact, several SNP-pairs were found correlated rendering the SNP-pair-based statistic significant. Possibly this may suggest several necessary compensatory mutations, or is reflecting passenger mutations. However, even many SNP-pairs collapse to only one molecular interaction. The higher predictive value obtained for miRNA-mRNA interactions compared to protein-protein pairs may reflect a more immediate need for compensatory mutations for interactions at the nucleotide level, whereas in proteins, synonymous vs. non-synonymous as well as the severity of the introduced amino acid change provide additional buffering of mutations. Taken together, while for actual prediction purposes, SNP-SNP correlation may not prove useful, a statistical signal, especially for miRNA-mRNA interactions, was clearly discernable. To put low predictive value in perspective, even in the case of within-protein correlated mutations using very distant sequences, and thus long evolutionary selection times, the accuracy of correlated mutations to indicate spatial contacts has been reported to be low (20%) (50, 51).

The above considerations assume the existence of a perfect gold standard. It is, however, possible that molecules do interact, but their interaction has not been observed or reported yet. And as we were able to demonstrate that pair hits are enriched on molecules known to interact, those that are found on molecules not (yet) reported to interact may, in fact, suggest that they may be functionally linked and may form physical contacts (Table 3), further supported by their increased semantic similarity (Figure 7). A large proportion of those putative interactions are of the type protein-protein interactions, which, of course, also reflects the genomic sequence space protein coding genes occupy in the Arabidopsis genome (Figure 1A). The predicted interactions await experimental confirmation or support by other data. Note that our method aimed to detect correlated mutations that have functional meaning and reflect functional associations, however, absence of correlated mutations does not mean absence of interaction. As with any correlative approach and as a general word of caution, high correlation levels cannot be taken as clear evidence of causality, but only to be in agreement with a causal relationship.

Pairs of tRNA-tRNA genes were found enriched for correlated mutations compared to intergenic regions (Figure 4). This may originate from tRNA-tRNA interactions in ribosomes as described by Smith and Yarus (52). The detected compensatory mutations may reflect contacts between the anticodon loops of the P-site and A-site of the tRNAs. Other RNAs such as miRNA and snoRNAs were also found enriched with regard to SNP-SNP correlations in comparison with intergenic regions. miRNA-miRNA correlations may perhaps be explained due to competitive mechanisms (53). Some RNA transcripts are able to communicate and co-regulate other RNAs by competing with its shared RNA regulator molecule, typically a miRNA (53–55). These are referred to as competing endogenous RNAs (ceRNAs) and the term “target mimicry” is often also used to describe this phenomenon in plants (56). In Arabidopsis, snoRNA are often polycistronic being transcribed from a single promoter and are also found to be dicistronic with tRNAs (57). The joint transcription suggests that those snoRNA and tRNA share regulatory mechanisms, transcriptional proteins and are evolutionary linked (58). Even though it is still obscure why these two types of molecules rely on the same transcriptional mechanisms, a joint functional role appears possible.

Of note also is the high number of novel interactions involving transposable elements (TEs). While the argument of SNP-density and covered genomic space of TE-genes holds as well, in the light of the reported functional roles of TEs, correlated mutations appear plausible as well. For example, it was shown that they act as gene expression regulators via small interfering RNAs (59) or by introducing novel cis-regulatory elements (60); i.e. different genomic sites are linked up, possibly reflected by correlated mutations.

Interestingly, pair hits were found enriched in gene loop contact regions and long-range chromosomal contacts (Figure 8) suggesting a link between 3D-genome organization and the sequence correlation structure. This may indicate that the integrity of such contacts is preserved in evolution via compensatory mutations. In Arabidopsis, different interacting elements for heterochromatin have been recognized. Heterochromatic islands and KNOT-engaged elements (KEEs) were found to be interacting (27, 28, 61). The contact regions of IHIs and KEEs present a high proportion of histone marks and TEs. The contact mechanism of these intra- and inter-chromosomal heterochromatic interactions remains unclear (62). Regions of high levels of chromatin interactions are rich in epigenetic marks such as DNA methylation, H3K9me2, and H3K27me1 (61, 63, 64). However, the notion of a critical role of DNA methylation or heterochromatin H3K9me3 marks to explain these contacts has been discarded, because in their absence, interactions remain the same (27). However, H3K27me3 was found to play a direct or indirect role in shaping the chromatin structure in plants (28, 62). These finding and the observed correlated mutation structure could suggest that TEs play a pivotal facilitator role in heterochromatin interactions.

TEs could be also be involved in short-range euchromatin interactions, so-called loops. In animals, euchromatin interacting modules have been designated Topologically Associating Domains (TADs). TADs-boundaries harbor binding sites for the CCCTC-binding factor (CTCF) that acts as an insulator, putting together two TADs-boundaries and forming a loop of the chromatin. Arabidopsis lacks CTCF to act as the insulator protein (65). However, over 1000 TADs-like regions were identified in Arabidopsis (28). The presence of these TAD-like regions suggests a similar mediator acting through unknown mechanisms (64). In Arabidopsis, regions have been found that seem to repel each other (28). Regions that are in the middle of the repelling regions are referred to as insulator-like regions. Thus, regions upstream and downstream of insulator-like regions have opposite interactions directionality. 400 insulator-like regions were recognized in Arabidopsis. As still little is known about CTCF-like proteins in plants, it is difficult to establish how relevant TEs are for these euchromatin interactions. However, in mammals, it has been found that species-specific TEs can influence chromatin interactions by modifying the binding sites of CTCF (66). The presented genome-wide and species-specific structure of correlated mutations may help to elucidate such specific mechanisms in plants by establishing links between genomic regions and the respectively encoded molecules.

In an attempt to identify particular molecule types or genomic elements enriched for correlated mutations, we compared all detected correlations on all possible such types to correlations found between intergenic regions (Figure 4). The rationale for this choice was the assumption that correlations between intergenic regions can be considered functionally neutral. However, and largely independent of the applied filtering, only about one-quarter of all possible pairs of molecules or genomic regions were found to exhibit more correlations than found between intergenic regions. Thus, correlation between intergenic regions may be more functionally relevant than thought and therefore occur more frequently than between many other types of genomic element pairs/molecules. Indeed, we observed an enrichment of intergenic-intergenic contacts in chromosomal regions exhibiting both physical contacts as well as showing elevated pair hit counts (Figure 8). Thus, the relevance of intergenic-intergenic contacts may lie in the maintenance of the global chromosomal 3D-architecture, which in turn may be reflected by an increased number of correlated mutations. And therefore, intergenic-intergenic contacts should not be considered neutral.

The continued and expanded sequencing efforts have yielded large SNP collections not only in Arabidopsis. The potential of these data to enable functional analyses has been demonstrated before, for example, specifically in *Arabidopsis thaliana* in the context of identifying novel cis-regulatory elements (67). Our study investigates and demonstrates the power of exploiting SNP information to unravel the global correlation structure of the Arabidopsis genome and to discern molecular interactions.

## Conclusions

By developing a series of filtering steps and random controls, our study was successful in establishing and applying the concept of correlated mutations to the identification of functional and physical interactions in closely related genomes within a single species by exploiting single nucleotide polymorphisms. This study succeeded in performing a genome-wide detection of the sequence-level correlation structure of a plant genome. As known physical interactions of molecules have been observed to be enriched for correlated mutations, correlated mutations on molecules not yet reported to interact or be functionally related may also have predictive value. With the rapidly increasing sequencing information, the approaches developed in this study may find broad application in other species.

## Acknowledgements

We wish to expressly thank Marco Ehlert (Max Planck of Colloids and Interfaces, Potsdam-Golm, Germany) for the critical help implementing an efficient computation strategy making the necessary massive computations possible. Further computing support was provided by Andreas Donath from Max Planck for Molecular Plant Physiology (Potsdam-Golm, Germany) and additional computational resources kindly made available by Alessandra Buonanno (Max Planck Institute for Gravitational Physics, Albert Einstein Institute, Potsdam-Golm, Germany), and the Max Planck Computer and Data Facility. We thank Ian Henderson for providing helpful comments.

## Declarations

### Authors’ contributions

DW conceived the study. LPJ performed all computations except for the co-expression analysis performed by DW. Both authors devised the strategies and methods of statistical testing, analyzed and interpreted the results, and wrote the manuscript.

### Competing interests

The authors declare that they have no competing interest associated with this study.

### Availability of data and materials

All information of data resources and methods necessary to replicate this study has been disclosed.

